# Extensive immune receptor repertoire diversity in disease-resistant rice landraces

**DOI:** 10.1101/2022.12.05.519081

**Authors:** Pierre Gladieux, Cock van Oosterhout, Sebastian Fairhead, Agathe Jouet, Diana Ortiz, Sebastien Ravel, Ram-Krishna Shrestha, Julien Frouin, Xiahong He, Youyong Zhu, Jean-Benoit Morel, Huichuan Huang, Thomas Kroj, Jonathan D G Jones

## Abstract

Plants have powerful defence mechanisms, and extensive immune receptor repertoires, yet crop monocultures are prone to epidemic diseases. Rice is susceptible to many diseases, such as rice blast caused by *Magnaporthe oryzae*. Varietal resistance of rice to blast relies on intracellular nucleotide binding, leucine-rich repeat (NLR) receptors that recognize specific pathogen molecules and trigger immune responses. In the Yuanyang terraces in south-west China, rice landraces rarely show severe losses to disease whereas commercial inbred lines show pronounced field susceptibility. Here, we investigate within-landrace NLR sequence diversity of nine rice landraces and eleven modern varieties of *indica, japonica* and *aus* using complexity reduction techniques. We find that NLRs display high sequence diversity in landraces, consistent with balancing selection, and that balancing selection at NLRs is more pervasive in landraces than modern varieties. Notably, modern varieties lack many ancient NLR haplotypes that are retained in some landraces. Our study emphasises the value of standing genetic variation that is maintained in farmer landraces as resource to make modern crops and agroecosystems less prone to disease.

## Introduction

Plant immunity requires timely activation of defence mechanisms, based upon detection of pathogen molecules via either cell-surface or intracellular immune receptors. Evasion of detection enables pathogens to proliferate and cause disease. When pathogens encounter large populations of genotypically identical and susceptible crop plants, rapid pathogen propagation and crop destruction can occur. *Resistance* (*R*) genes usually encode intracellular NLR (nucleotide binding, leucine-rich repeat) immune receptors which detect specific pathogen effectors (virulence factors) and confer the innate ability to recognise pathogens. Most plants carry hundreds of NLR-encoding genes^1^ and display extensive variation at *R* gene loci. In host-parasite coevolution, extensive standing variation at these *R*-genes is critical to cope with evolutionary diversity in pathogens, which enables sustainable resistance in natural host populations^2^.

Genetic diversity in hosts for pathogen recognition can slow epidemics. Variation between host genotypes in their resistance to different pathogen strains reduces the risk that the host population is overcome by a single pathogen strain^3,4,5^. Importantly, population-level resistance can be thought of as an emergent property resulting from diversity in immune receptor repertoires, and a single “perfect” genotype cannot capture this property. In contrast, standard plant breeding practice in modern agriculture requires varieties to display uniformity and reliable performance over a wide range of environments. Such properties of modern agroecosystems are incompatible with population-level heterogeneity in immune receptor repertoires. Traditional farming systems tend to rely on genetically heterogeneous mixtures of traditional varieties, referred to as landraces^6^, and they often provide effective and sustainable disease control^7^. For example, traditional farmers in the Yuanyang terraces in Yunnan (south-west China) cultivate rice landraces that rarely show severe losses to infectious diseases^8,9^.

About 200 landraces^10^ are maintained by a traditional social organization involving sporadic seeds exchange between farmers^11^. Furthermore, farmers subconsciously carry out varietal selection by not planting varieties that were heavily impacted by disease in the previous season^11^. This social organization may have contributed to *R* gene heterogeneity in two ways: (1) by enhancing spatiotemporal variation in *R* gene repertoires and intensifying selection through selective planting, and (2) by increasing gene flow (through the exchange of seeds). Both processes may have contributed to resistance durability in rice populations grown in the Yuanyang terraces.

Modern farming practices have profound coevolutionary implications, and these can be best understood in the light of population genetic theory^12-15^. During coevolution, adaptations in one species provoke counter-adaptations in the coevolving species. Consequently, the direction and intensity of natural selection constantly change^16^. Assuming that both antagonists possess sufficient genetic variation to fuel these continuous adaptations, none of the interacting species gains a sustained fitness advantage. Balancing selection maintains genetic polymorphisms in host resistance genes due to spatiotemporal variation in selection pressures posed by the pathogens. In other words, different genetic variants (e.g., alleles or haplotypes) are favoured in different places and different times, meaning that genetic polymorphism can be maintained long-term. This is known as the trench-warfare model^17^. Importantly, this also limits the infection incidence (i.e., the number of infected hosts), because the susceptible host genotype is locally and/or temporally continuously replaced by a genotype that is resistant to the prevailing pathogen strain. The composition of the prevailing pathogen strains is itself variable. This makes antagonistic coevolution a zero-sum game with no knockout winners or losers.

In contrast, if there is insufficient host genetic variation, a pathogen strain that can overcome the defences of the predominant host genotype is likely to cause damaging reductions in host fitness if the population lacks any resistant host plants. If the host population survives, the susceptible genotype may be lost completely (because of the unrestrained, exponential increase of the winning pathogen strain). In turn, this tends to result in a turnover of sequence variation, and this type of host-parasite coevolution, which matches the experience of plant breeders releasing new varieties that are monocultures, is known as the arms-race model^17^. Modern crops consisting of genetically near-uniform host plants are ill-equipped to face the co-evolutionary challenges posed by diverse, rapidly evolving pathogens with a trench-warfare model. Rather, they are forced into an arms-race that requires a continuous input of novel resistant varieties developed by plant breeders (as well as agrichemical disease-control measures) to keep pace with their rapidly evolving pathogens. In other words, the standing variation implied by the trench-warfare model is a “recycle- and-reuse” strategy that is sustainable, whereas the arms-race model uses sequence variation in a disposable fashion, making it less sustainable. In this study, we examine this coevolutionary hypothesis (see ref. ^2^ for an excellent review).

We report here on a study to examine whether *indica* landraces of the Yuanyang terraces might show relatively elevated levels of diversity in their NLR immune gene repertoires, hypothesising that traditional farming practices are better than modern breeding at conserving variation. We use RenSeq sequence capture^18^ to enrich NLR sequences prior to sequencing, and we designed a set of biotinylated bait sequences to capture NLR-encoding immune receptors, based on reference genomes of *O. sativa ssp japonica* and *O. sativa ssp indica* (herein referred to as japonica and indica, respectively). All rice genotypes were assessed using Illumina sequencing of captured NLRs. We evaluated RenSeq data from 11 japonica, aus, and indica inbreds, and we compared these data to those from 38 accessions from seven different landraces from the Yuanyang terraces. We evaluated presence/absence variation and sequence diversity. These analyses revealed a marked depletion of NLR polymorphisms in the japonica and indica inbred lines and substantial within-landrace NLR sequence heterogeneity that likely underpins the relatively low incidence of rice blast in Yuanyang terraces^19^. We discuss the demographic events and evolutionary forces that can explain these data.

## Results

### Data generation

We selected 49 Asian rice (*Oryza sativa*) accessions for RenSeq analysis, representing seven indica landraces (36 accessions), two japonica landraces (two accessions), and eleven modern varieties of indica, japonica, and aus (11 accessions). Landrace accessions were sampled in 2014 and 2015 in the fields of traditional rice farmers in three villages from the Yuanyang rice terrace region in Yunnan. RenSeq baits were designed to hybridize with 761 NLR-coding sequences from japonica and indica rice. To generate a baseline against which to identify features of polymorphism in genes of interest, 68 accessions were also characterized using genotyping-by-sequencing (GBS), representing nine landraces and 28 modern varieties (Supplementary Table 1). After standardizing the number of reads, read-mapping, SNP calling, and masking of paralogous calls or other SNPs with excess heterozygosity, the RenSeq dataset included 40,530 biallelic SNPs (reference sequence: 596 genes, 2.4 Mb) and the GBS dataset 199,130 biallelic SNPs (reference sequence: 42,031 genes, 99.6 Mb).

### Population subdivision

To understand the genetic relationships among rice subspecies and landraces, and to investigate signatures of natural selection at the intraspecific level, we inferred population structure from RenSeq and GBS data using complementary approaches that make no assumption about Hardy-Weinberg equilibrium and are therefore appropriate to analyse structured or inbred populations. Both clustering analyses with the SNMF software^20^ (Supplementary Figure 1) and neighbor-net phylogenetic networks^21^ (Figure 1) revealed consistent patterns that split genetic variation primarily by type of rice: aus, modern temperate japonica, modern tropical japonica, modern indica, indica landraces, japonica landraces.

**Figure 1.**
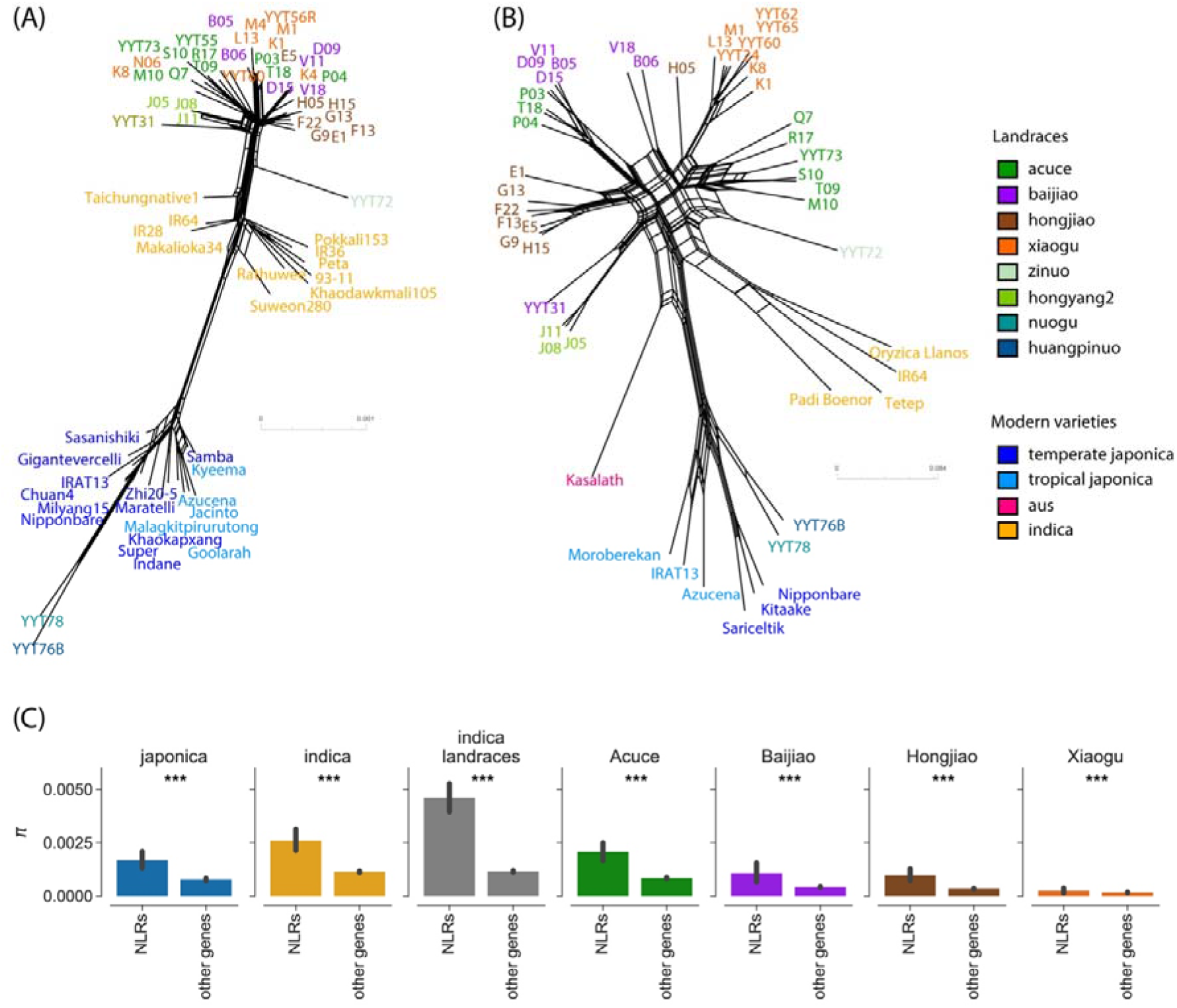
Analysis of nucleotide diversity separates modern varieties and landraces, and reveals high standing genetic diversity in landraces compared to modern varieties. Population subdivision was inferred from 49 and 68 accessions, for GBS (A) and RenSeq (B), respectively, representing 13 varieties or landraces shown with different colors. Neighbor-net phylogenetic networks estimated with Splitstree 21 for GBS and RenSeq data. Reticulations in the network indicate phylogenetic conflicts caused by homoplasy. Splitstree analysis was based on 31,770 biallelic SNPs with less than 80% missing data for RenSeq data, and 60,166 biallelic SNPs with less than 50% missing data for GBS data. Panel (C) represents bar plots of nucleotide diversity π in RenSeq data (NLRs) and GBS data (other genes). The ‘indica landraces’ group includes one randomly chosen accession per individual landrace; one out of 30 resamples of one accession per landrace is included in the plot, and summary statistics for the remaining 29 resamples are presented in Supplementary Table 2. Error bars represent the standard error. Comparisons between NLRs and other genes: **p<0.01, ***p<0.001 (Mann-Whitney U-tests). Comparisons between indica or japonica and the group of indica landraces for NLRs and other genes were all significant (Mann-Whitney U-tests in Supplementary Figure 2 and Supplementary Figure 7).

Within indica landraces, accessions from the same variety tended to cluster together, except for three Acuce accessions that clustered with Baijiao accessions, and one Baijiao accession that clustered with Hongyang2 (Figure 1; Supplementary Figure 1). Note that by plotting the two networks in the same scale, the GBS network would be much more compact than the RenSeq network. This indicates that the level of differentiation is higher at NLRs compared to the remainder of the genome, consistent with directional (positive) selection on the NLRs.

Clustering analyses further revealed that Acuce accessions V18 and B06, and Hongjiao accession H05, had mixed ancestry in multiple clusters at most K values (Supplementary Figure 1), and did not branch with other accessions from the same varieties in the neighbor-net network (Figure 1). These accessions likely represent genetically introgressed (hybrid) lineages.

### Demography of modern varieties and landraces

We used GBS sequences in non-NLR genes to explore how population history shaped patterns of genome-wide polymorphism in the different rice populations. Nucleotide diversity (π) differed significantly between populations (Kruskal-Wallis test: H=11330.1, d.f.=6, p <0.001), and most post-hoc pairwise comparisons were statistically significant (p<0.001; Mann-Whitney tests with Bonferroni-Holm correction; Supplementary Figure 2). Seven indica landrace accessions harbored almost as much nucleotide diversity (from π=0.00089/bp to π=0.00108/bp across 30 random resamples including only one accession per population) as eleven modern indica varieties (π=0.00107/bp), and more nucleotide diversity than seventeen modern japonica varieties (π=0.00060/bp; Figure 1C; Supplementary Figure 2; Supplementary Table 2; Supplementary Figure 3). Nucleotide diversity in individual landraces ranged from π=0.00018/bp in Xiaogu to π=0.00075/bp in Acuce (Figure 1C; Supplementary Figure 2; Supplementary Table 2; Supplementary Figure 3). These patterns of polymorphism indicate that farming practices have maintained a relatively high level of genome-wide standing variation in landraces.

The frequency distribution of polymorphisms, as measured by Tajima’s D, indicated that modern landraces and varieties have experienced distinct evolutionary and/or demographic processes. The average D across GBS loci was close to zero across all indica landraces as a group (from D=-0.046 to D=0.249 across 30 random resamples including only one accession per population), indicating mutation-drift equilibrium. In other words, there is no evidence of selection or significant demographic changes across the indica landraces. In contrast, at the scale of individual landraces, the average D indicated an excess of low frequency variants in Acuce (D=-0.339), Baijiao (D=-0.242), Xiaogu (D=-0.095), and a shift toward higher frequency alleles in Hongjiao (D=0.686) and in the two populations of modern varieties (indica: D=0.444; japonica: D=0.473) (Supplementary Table 2; Supplementary Figure 2). This indicates that the individual landraces have followed distinct demographic histories and/or selection pressures and represent distinct populations with unique evolutionary histories.

To more accurately estimate the demographic history of the different rice groups, we used coalescent simulations within an Approximate Bayesian Computations (ABC) framework^22^ to compare demographic models. Posterior probabilities supported different demographic models for individual landraces. The best-supported models were an exponential growth model for Acuce and Baijiao, a bottleneck model for Xiaogu, and a two epochs model with population contraction for Hongjiao (Supplementary Table 3; Supplementary Figure 4). For indica landraces as a group, the constant size model was the best supported, while two epochs model with population contraction had higher posterior probabilities for indica and japonica. Different landraces had different population dynamics, likely due to changes in their popularity with farmers. Nevertheless, when considering the landraces as a single metapopulation, the demography of rice in the Yuanyang terraces remained constant, with temporal changes experienced by distinct landraces cancelling each other out. This suggests that the traditional agroecosystem kept the total metapopulation size relatively constant over centuries.

The area occupied by modern varieties is much larger than the terraces of Yuanyang, which implies that the modern varieties have a larger census population size than all landraces combined. However, our demographic modelling points to a decreasing population size, which reflects the fact that modern agroecosystems are based on the cultivation of a limited number of related genotypes. Despite their larger census population size, the modern varieties are likely to experience more genetic drift and/or selective sweeps than the landraces, which exposes modern varieties to genomic erosion^23^. In the following, we use the demographic histories inferred from GBS loci as baselines to test for selection at NLRs.

### Linking NLR diversity to function and presence/absence variation

Before testing the impact of selection at NLRs, we first examined the factors accounting for the molecular variability of NLRs. To test whether variation was evenly distributed among the different protein domains of NLRs, we used interpro to define domains and computed summary statistics at the scale of domains. Nucleotide diversity (π) and the ratio of non-synonymous to synonymous nucleotide diversity (πN/πS) differed significantly between protein domains both at the species-wide scale (Kruskal-Wallis test: H=18.4, d.f.=3, p=0.001 for π; H=20.4, d.f.=3, p=0.0004 for πN/πS; Figure 2) and at the scale of individuals varieties and landraces (Supplementary Table 4; Supplementary Figure 5). Nucleotide diversity at the leucine-reach repeats (LRR) was significantly higher than nucleotide diversity at the coiled-coil (CC), and nucleotide-binding (NBARC^24^) domains (Post-hoc Mann-Whitney test: p=0.016 and p =0.037, respectively; Figure 2A), and the πN/πS at the LRR was significantly higher than πN/πS at the CC domain (Post-hoc Mann-Whitney test: p=0.0006; Figure 2A).

**Figure 2.**
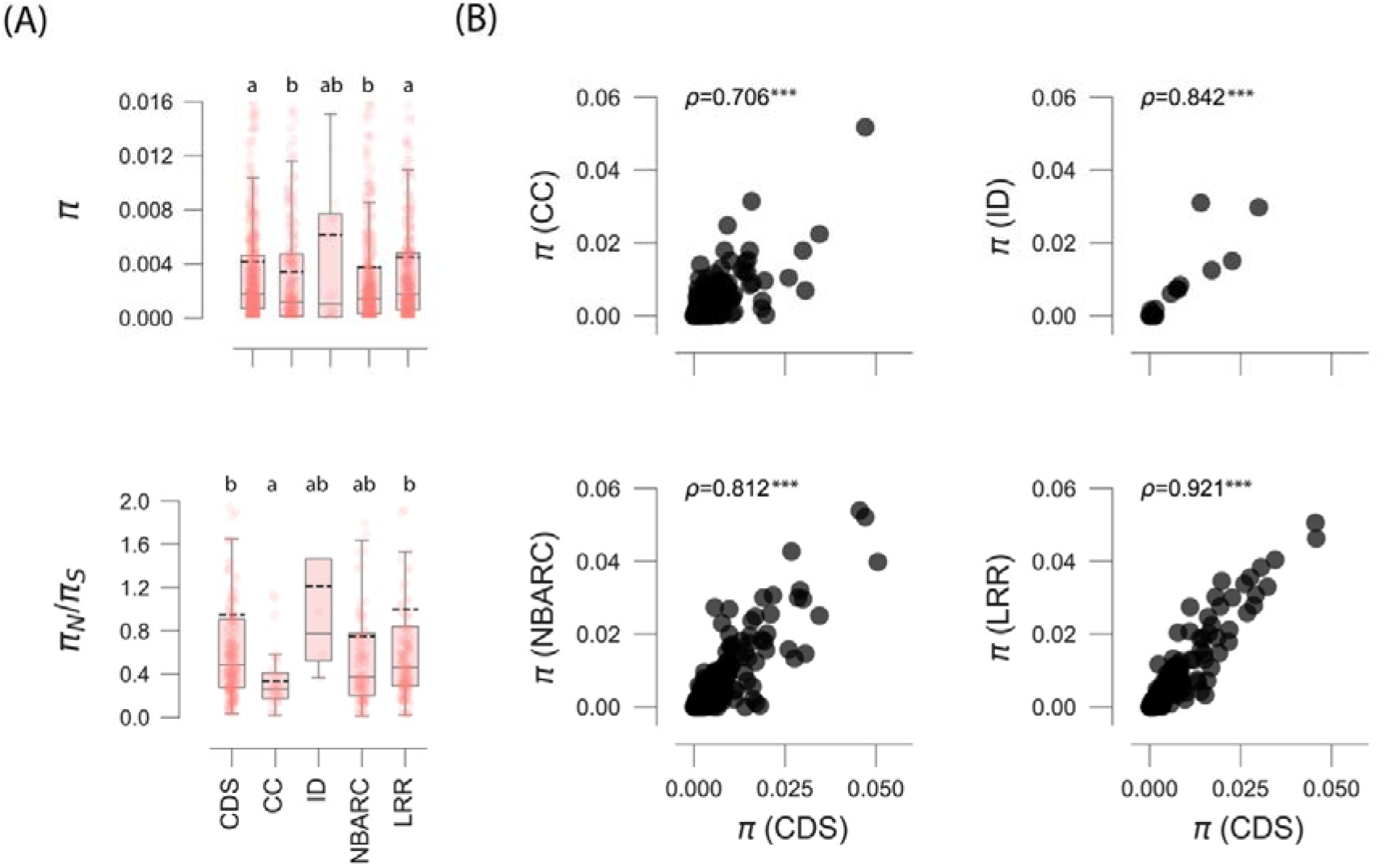
Patterns of nucleotide variation in NLR genes. (A) Species-wide nucleotide diversity (π) and ratio of non-synonymous to synonymous nucleotide diversity (πN/πS) in full coding sequence (CDS) and functional domains (CC: coiled-coil; ID: integrated domain^40^; NBARC: nucleotide-binding domain^24^; LRR: leucine-rich repeats). (B) Correlation between nucleotide diversity in domains and full coding sequences. In panel (A), a number of data points were cropped from the nucleotide diversity plot for visually optimal presentation, but included in statistical tests. In boxplots, dashed black line is mean, solid black line is median. The letters a and b above the boxplots indicate whether the distributions are similar (when sharing the same letter), or significantly (p<0.05) different, based on a Mann-Whitney test. In panel (B), ρ is Spearman’s rank correlation coefficient (***p<0.0001)

Nucleotide diversity in coding sequence was most strongly correlated with nucleotide diversity in LRR compared to nucleotide diversity in the CC or NBARC domain (Figure 2B). We conclude that LRR variation is the best predictor of NLR molecular diversity, consistent with the central role of the LRR domain in recognition, and thus in trench warfare coevolution with cognate ligands.

We used normalized read mapping depth to investigate the impact of presence/absence variation on the molecular variability of NLRs. At the species level, we found significant positive correlations between presence/absence diversity and nucleotide diversity (Spearman’s rank correlation coefficient ρ=0.25, p<0.001) (Figure 3A). Species-wide nucleotide diversity π was significantly higher in core NLRs compared to accessory NLRs (π=0.00455 for core NLRs, π=0.00338 for accessory NLRs; core NLRs are present in all accessions of all subsamples of two accessions from a given population; Figure 3B), and the same pattern was observed at the population level except in Xiaogu (Mann-Whitney U tests, p<0.0001; Supplementary Figure 6). The maintenance of greater nucleotide diversity in core NLRs compared to accessory NLRs suggests stronger balancing selection could be acting on core NLRs.

**Figure 3.**
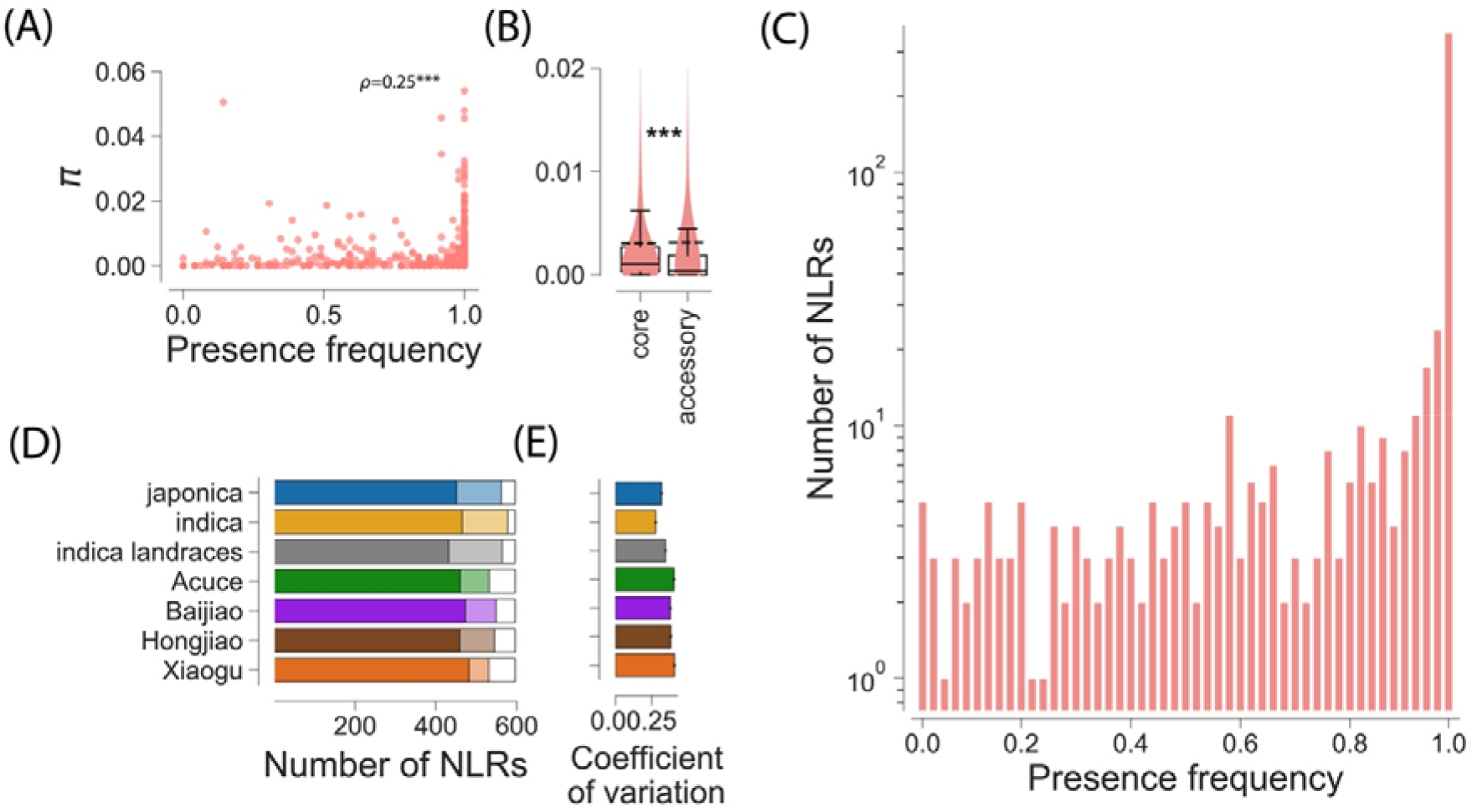
Presence-absence variation of 596 NLRs in 49 rice accessions. (A) Species-wide nucleotide diversity (π per bp) vs. presence frequency of NLRs; ρ is Spearman’s rank correlation coefficient (***p<0.001). (B) Species-wide nucleotide diversity (π per bp) in core and accessory NLRs, ***p<0.001; Mann-Whitney posthoc test with Holm-Bonferroni correction; in box-plots dashed black line is mean, solid black line is median. (C) Distribution of NLR presence frequency. (D) numbers of core (dark), accessory (light) and missing (white) NLRs, a core (missing) NLRs being present (absent) in all accessions of all subsamples of two accessions from a given population. (E) Jackknife estimates of coefficient of variation in number of NLRs present, with error bars representing confidence intervals. The ‘indica landraces’ group includes one randomly chosen accession per individual landrace; one out of 30 resamples of one accession per landrace is included in the plot.

NLRs showed remarkable levels of presence/absence variation. Approximately 50% of the NLRs (358 NLRs out of the 596 NLRs used as reference sequences for read mapping) were present in all accessions of all populations of modern varieties and landraces, and these can be considered species-level core-NLRs. Of the remainder, ca. 30% were present in less than 90% of all accessions (Figure 3C). At the population level, the number of core-NLRs was similar in modern varieties (451 in japonica, 465 in indica) and in landraces (460 in Acuce, 482 in Xiaogu, 473 in Baijiao, 459 in Hongjiao; Figure 3D), and most NLRs that were core in a given population were core in all populations (Supplementary Figure 6).

Interestingly, though, the variation in number of NLRs per population was higher in landraces than in modern varieties (Figure 3E). Presence/absence variation in NLR repertoires was significantly higher for landraces from different populations (median = 73) than for modern varieties of different japonica types (median = 65), (Mann-Whitney, W = 513434.0, p=0.0104). In other words, two randomly picked plants from two landraces differ more in their NLR repertoire than two randomly picked plants from temperate or tropical rice populations. All NLRs from the chromosome 7 of the indica reference genome were missing in all accessions of Acuce and Xiaogu (Supplementary Figure 6). Population-level presence frequency distributions followed the same reversed L-shaped distribution at the species-wide level (Supplementary Figure 6).

### Impact of balancing selection on overall NLR variation

Comparisons of nucleotide diversity (π) at NLRs between landraces and varieties further revealed statistically significant differences between indica modern varieties and individual landraces, as well as between indica modern varieties and indica landraces as a group (i.e., measured using only one seed per bag of seeds) (Kruskal-Wallis test: H=814.1, d.f.=6, p<0.0001; Mann-Whitney post-hoc tests in Supplementary Figure 7). Individual landraces (“bag of seeds”) harbored 7% (in Xiaogu) to 56% (in Acuce) of the total nucleotide diversity measured in modern indica. Even more remarkably, a single “bag of seeds” of Xiaogu and Acuce contained 17% to 134%, respectively, of the nucleotide diversity measured in all modern japonica. Seven indica landraces displayed similar or significantly higher nucleotide diversity (from π=0.00343/bp to π=0.00436/bp, across 30 independent resamplings of one accession per “bag of seeds”) than four modern indica (π=0.00343/bp) and six modern japonica (π=0.00144/bp; Figure 2; Supplementary Table 2; Supplementary Figure 7). The observed differences in nucleotide diversity indicate that traditional breeding of landraces maintained higher molecular diversity at immune receptors than breeding and improvement of modern varieties. This result thus corroborates the demographic analysis, showing that modern varieties experienced more genomic erosion than the landraces. Given that the census population size is likely to be larger for the modern varieties, the difference in genomic erosion is likely to be the result of differences in selection pressures. In particular, modern varieties may have experienced more intense directional selection (potentially resulting in selective sweeps), and/or conversely, landraces may have experienced more balancing selection that maintained diversity at their NLRs.

To test for balancing selection at NLRs as a group, we corrected for the deviation from the standard demographic equilibrium by simulating datasets of similar number of sequences and loci as the observed NLR datasets according to the best supported demographic models estimated from GBS data. Nucleotide diversity π observed in RenSeq data was higher than expected in all populations (p<0.0001) and Tajima’s D was higher than expected in all populations (p<0.0001) except Xiaogu and Hongjiao (p=0.115 and p=0.293, respectively; Supplementary Figure 8). Higher nucleotide diversity at NLRs was also supported by comparisons between GBS sequences from NLR and non-NLR genes (Supplementary Table 5). Assuming that the mutation rate is not elevated at NLRs, these findings indicate that in both modern varieties and landraces, balancing selection has maintained molecular variation at NLRs. This analysis thus rules out that directional selection (or selective sweeps) has resulted in more severe genomic erosion of NLRs in modern varieties. (Note that this does not preclude the possibility that directional selection in modern varieties may have eroded genetic diversity elsewhere in their genome). Next, we examined whether landraces may have experienced more balancing selection than modern varieties, which is an alternative hypothesis that could explain their relatively elevated NLR diversity.

### Search for NLRs under balancing selection

To identify NLRs under balancing selection, we mapped the observed values of nucleotide diversity π and Tajima’s D of each NLR on the joint density of (π, D) expected under selective neutrality while accounting for the demographic history of each group. NLRs were identified as under balancing selection if falling in the top 5% of π and D values calculated on datasets simulated under the best supported demographic models. Five and 19 NLRs were identified as being under balancing selection in modern indica and japonica, respectively (Supplementary Table 6). Across all individual landraces, on average 35.3 NLRs were identified to be under balancing selection in 30 resamplings of one accession per landrace (standard deviation: 3.3; range: 31 – 41; Figure 4A; Supplementary Table 6). This analysis conclusively shows that overall, NLRs in landraces appear to be under stronger balancing selection compared to NLRs in modern varieties, and this could explain why the NLR diversity of landraces is so highly elevated.

**Figure 4.**
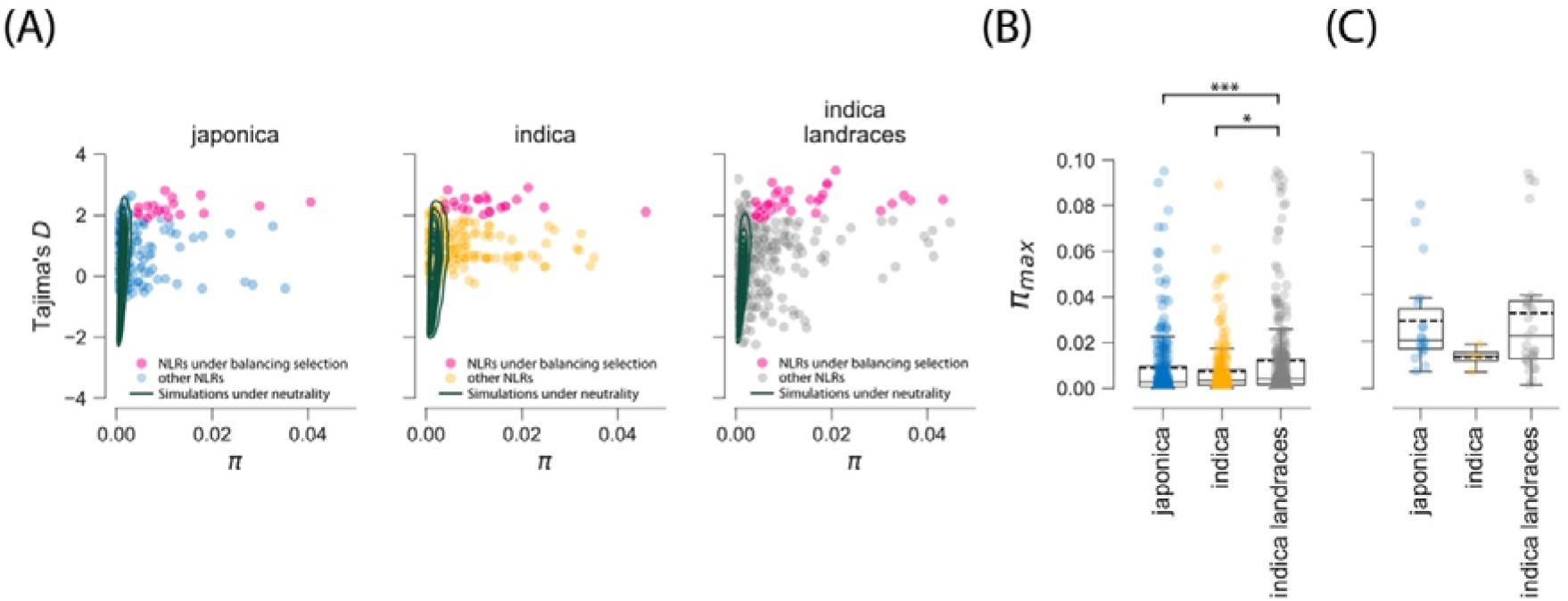
NLRs in indica landraces display signatures of balancing selection and enrichment in long-lived alleles. (A) Tajima’s D and nucleotide diversity π in NLRs from modern japonica varieties, modern indica varieties and indica landraces; lines represent kernel density estimates of summary statistics computed on 1000 datasets simulated for each NLRs, with datasets being of the same sample size and sequence length as NLR sequences; simulations were carried out by sampling multivariate parameters from posterior distributions of the best supported demographic models; (B) π_max_ in modern indica varieties and indica landraces, computed for all NLRs; (C) π_max_ in modern indica varieties and indica landraces, computed for all NLRs under balancing selection. Kernel density estimate plots were produced using the Python package Seaborn 0.11.2. π_max_ represents the maximum number of pairwise differences and measures the maximum depth of gene trees. *p<0.05, ***p<0.001, one-sided Mann-Whitney tests with Bonferroni-Holm correction. The ‘indica landraces’ group includes one randomly chosen accession per individual landrace; one out of 30 resamples of one accession per landrace is included in the plot.

Fourteen NLRs deviated from selective neutrality in all 30 resamplings. Chromosome 11 harbored most of the NLRs under balancing selection, with one NLR in indica, five NLRs in japonica and five to eleven NLRs in 30 resamplings of indica landraces (Supplementary Figure 9. Three NLRs (BGIOSGA022757, BGIOSGA024307, BGIOSGA033570) were under balancing selection in both japonica and indica landraces and one NLR (BGIOSGA028563) was under balancing selection in both indica and indica landraces (Supplementary Table 6).

π_max_, which represents the maximum number of pairwise differences and measures the maximum depth of gene genealogies, was significantly higher in all 30 resamplings of indica landraces than modern indica and japonica varieties (one-sided Mann-Whitney U tests; p<0.05; Figure 4B), indicating that indica landraces have kept older NLR alleles than modern varieties. For instance, unlike landraces, modern indica varieties lacked NLRs with π_max_ in the range 0.089-0.145, which corresponds to a minimal allelic divergence of T=6.8 million years (assuming π_max_=2µT, with µ=6.5e-9/bp^25^). NLRs had deeper genealogies in indica landraces (across 30 resamplings: average π_max_: 0.0317-0.0319, median π_max_: 0.0204-0.0225, max π_max_: 0.103) than in modern indica or japonica varieties, which showed more recent common ancestry of alleles (in indica, average π_max_: 0.013, median π_max_: 0.014; max π_max_: 0.019; in japonica, average π_max_: 0.029, median π_max_: 0.020, max π_max_: 0.078; Figure 4C). These patterns indicate that, compared with landraces, ancient NLR polymorphisms have been lost due to more severe genomic erosion in modern varieties.

Among NLRs under balancing selection in indica landraces featured gene *RGA4* (*BGIOSGA034263*), which is involved in resistance to rice blast. *RGA4* was under balancing selection in 22 out of 30 resamplings of one accession per landrace, and displayed relatively high values of nucleotide diversity π and Tajima’s D in indica landraces (across 30 resamplings: average π= 0.0062, standard deviation π=0.0005; percentile range π=[69.7%-80.4%], D=1.925, standard deviation D=0.51, percentile range D=[31.9%-97.0%]) as well as in modern indica (π=0.0051, percentile(π)=79.1%, D=0.887, percentile(D)=65.1%) and japonica varieties (π=0.0053, percentile(π)=82.9%, D=0.302, percentile(D)=40.2%).

Maximum-likelihood gene genealogy revealed standing variation exclusive to the landraces at *RGA4* (Figure 5A). The high molecular diversity detected at *RGA4* was mostly driven by the LRR domain in indica landraces, and the LRR and NBARC domain in modern indica varieties (Figure 5B). In addition to *RGA4*, five other NLRs, out of the 32 NLRs with a signature of balancing selection in indica landraces, were involved in head-to-head pairs of NLRs, which is more than expected by chance (Fisher exact test, p=0.0014). More generally, paired NLRs harbored significantly more nucleotide diversity, and more anciently diverged alleles than other NLRs in indica landraces (Supplementary Figure 10; one-sided Mann-Whitney tests with Bonferroni-Holm corrections; π: H=3063.0, p=0.030; π_max_: H=4727.0, p=0.006). Nucleotide diversity values within head-to-head pairs were also correlated (π: Spearman’s ρ=0.76, p=0.005) (Supplementary Figure 10). Such signatures of coevolution were not observed for the *RGA4*/*RGA5* pair. RGA5, the NLR which binds to effectors AVR-Pia and AVR-CO39, was polymorphic at the species level, but monomorphic in indica landraces and modern japonica and indica. *RGA5* was also less frequent than *RGA4*: while *RGA4* was detected in all 49 accessions, *RGA4* was present in only 22.

**Figure 5.**
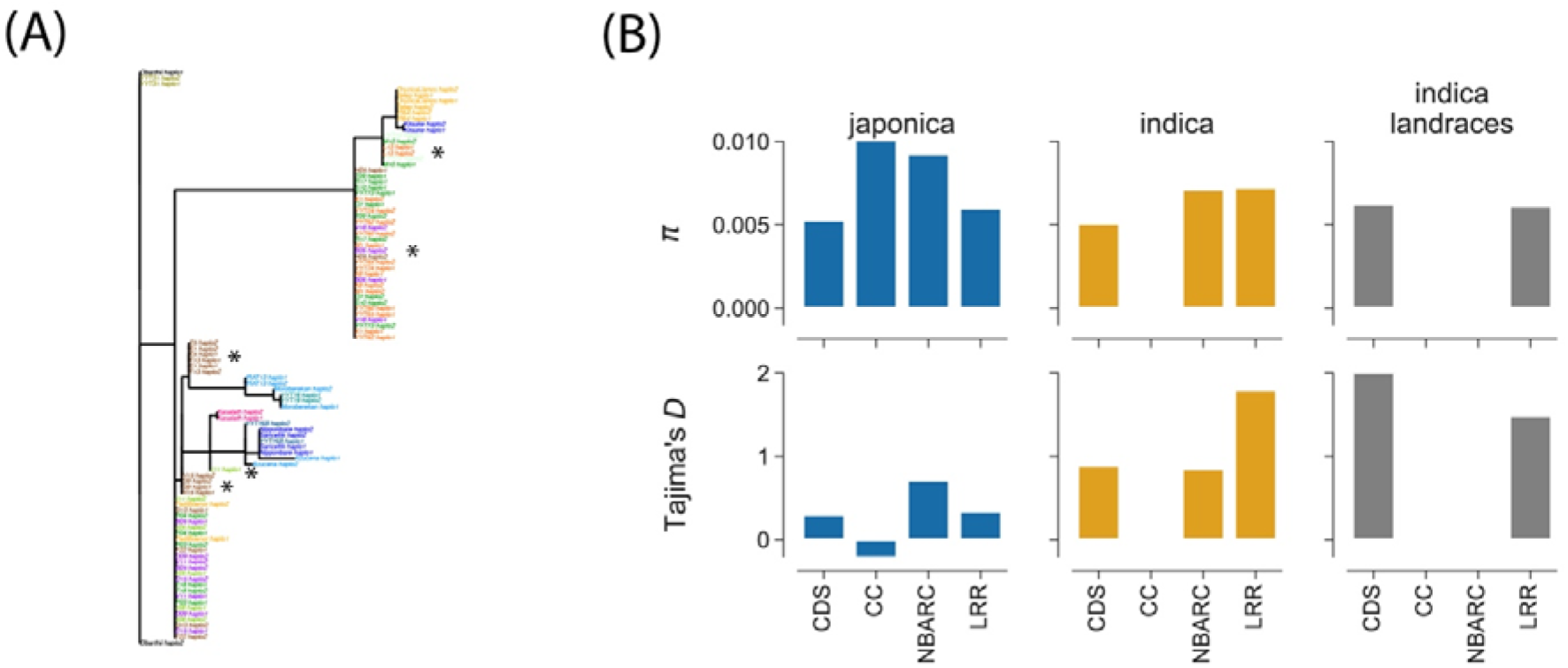
Balancing selection at RGA4. (A) Maximum likelihood phylogenies. Stars indicate haplotypes exclusive to landraces (B) Nucleotide diversity π and Tajima’s D in full coding sequence (CDS) and functional domains. The ‘indica landraces’ group includes one randomly chosen accession per individual landrace; one out of 30 resamples of one accession per landrace is included in the plot.

### Impact of recurrent directional selection on NLR variation

Pathogen-mediated balancing selection is likely to be ancient, which is suggested by the observed deep gene genealogies of NLRs. To examine this further, we identified NLRs with signatures of adaptation acting over longer time periods in modern varieties and landraces. We therefore used a Bayesian extension of the McDonald-Kreitman test implemented in the SnIPRE program^26^. In particular, we compared polymorphism and divergence at synonymous and non-synonymous sites in NLRs for which an orthologous sequence could be identified in the outgroup *O. barthii*. In both indica landraces and modern indica varieties, we detected widespread purifying selection against strongly deleterious mutations in almost all NLRs.

Purifying selection thus reduces the genetic load in almost all NLRs, indicating that their nucleotide sequence is functionally constrained. Nevertheless, between one to four NLRs from the 285 polymorphic NLRs with outgroup data showed evidence of directional selection across the 30 resampled groups of indica landraces (Supplementary Table 7; Supplementary Figure 11). Three NLRs (*BGIOSGA027982, BGIOSGA040540, BGIOSGA024574*) showed a consistent signature of directional selection, being flagged up in >14 of the 30 resampled groups of indica landraces (Figure 6). In contrast, none of the NLRs displayed significant directional selection in modern indica and japonica (Figure 6). Apparently, some NLRs show evidence of adaptive evolutionary change, possibly in response to changes in pathogen pressures, but this signature is only observed in indica landraces, not in modern indica and japonica.

**Figure 6.**
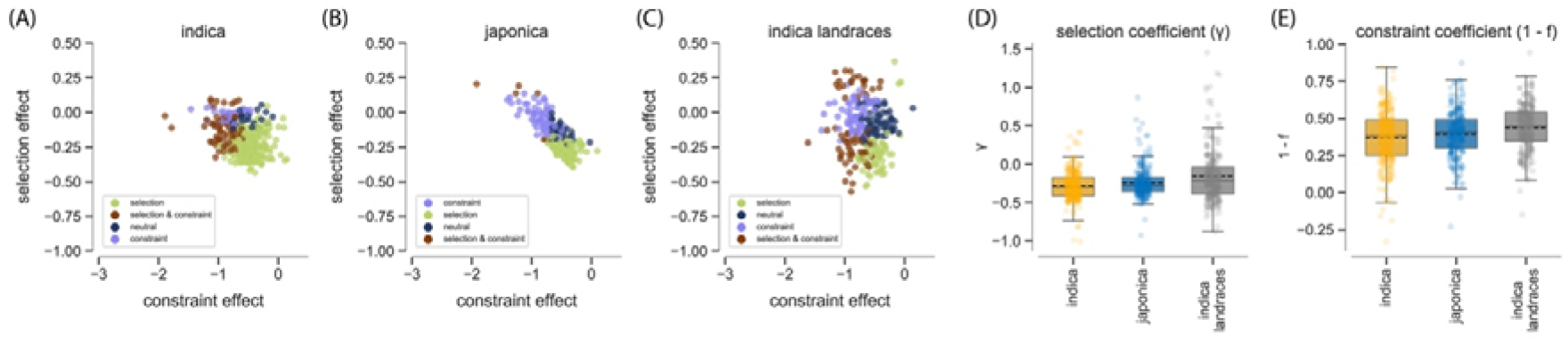
SnIPRE estimates of recurrent directional selection in 285 NLRs with outgroup data. (A) Selection and constraint effects in indica, (B) Selection and constraint effects in japonica, (C) Selection and constraint effects in indica landraces, (D) Selection coefficients in indica, japonica and indica landraces, (E) Constraint coefficients in indica, japonica and indica landraces. The selection effects reflect the selection coefficients (γ), with γ>0 indicating positive selection and γ<0 negative selection. The constraint (or non-synonymous) effects reflect mutational constraint (1-f, f being the proportion of non-synonymous mutations that are not lethal). The ‘indica landraces’ group includes one randomly chosen accession per individual landrace; one out of 30 resamples of one accession per landrace is included in the plot, and results for the remaining 29 resamples are presented in Supplementary Figure 11

Differences between the landraces and modern varieties were also revealed when analysing and comparing the selection coefficient γ and the constraint coefficient 1 – f using the SnIPRE program. Both statistics were significantly different between indica landraces and modern indica varieties in 28 and 30 resampled indica landraces groups, respectively (Mann-Whitney U tests, p<0.05; Supplementary Table 7). Furthermore, the mean values of both coefficients were greater in indica landraces than modern indica varieties for all 30 resampled groups (Figure 6; Supplementary Figure 11). Again, these analyses suggest that more adaptive evolutionary changes are occurring in the indica landraces than in the modern varieties. Our data are consistent with the hypothesis that NLR diversity plays a role in population-level resistance against rice pathogens, and suggest that landraces also provide a rich source of additional recognition capacities that could be recruited into modern varieties.

## Discussion

We used a combination of RenSeq sequence capture and Illumina sequencing to provide a comprehensive overview of nucleotide polymorphism at NLR-encoding loci in 49 accessions of *O. sativa*. In all modern and landrace populations, nucleotide diversity at NLRs was consistently higher than at other loci characterized using genome complexity reduction sequencing, and significantly higher than predicted from models of neutral evolution fitted to the other loci. NLRs thus appear to be highly variable in plant genomes at the intraspecific level, similar to other types of immune receptors outside the kingdom Plantae^27-30^. The high diversity of NLRs is consistent with their involvement in coevolutionary interactions with pathogen-derived ligands that impose strong selection on NLRs^17,31^. Pathogen-mediated selection can result in balancing selection (which maintains diversity), and/or directional selection (which results in changes in variation). If directional selection changes across time and space, for example due to changes in the composition of local pathogen communities, directional selection can act like balancing selection, and help to maintain diversity ^16^.

Fundamentally, whether selection is directional or balancing determines the mode of coevolution; it leads to a Red Queen arms race with tit-for-tat changes and a transient polymorphism, or it may result in a trench warfare model with a long-lasting standing variation and balanced polymorphisms^17^.

Not all NLRs are hypervariable, and the observed range in patterns of diversity included ∼20% loci without polymorphism. Lack of diversity might reflect the fact that some NLRs can contribute to downstream signaling^32^, which may impose strong purifying selection to maintain the function. However, no helper NLRs have been functionally defined in grasses. Some NLRs such as ZAR1 have a broadly conserved role in detecting effector manipulation of protein kinases involved in signal transduction. We are skeptical that the number of rice NLRs we observed that show little diversity can be thus explained. Strong directional selection (e.g., by a ubiquitous, dominant pathogen) can also reduce variation by fixing a (likely temporarily) selectively favoured allele. The correlation between the diversity of the coding sequence of NLRs with the diversity of their LRR domains suggests that the main driver of the diversification of NLRs is the selective pressure exerted on this domain, and therefore on the recognitional capacity of the NLR.

Another important observation is that indica landraces carry significantly more nucleotide diversity at NLRs than modern indica and japonica varieties. Similarly, the presence/absence variation in number of NLRs per population was also higher in landraces than in modern varieties. The high diversity is not only observed at the scale of the agroecosystem, but it is also observed within landraces; there is as much nucleotide diversity in nine individual lines from the ‘Acuce’ accession as in six japonica modern varieties. In other words, nine individual plants of a landrace may possess as much NLR diversity as what could be found among billions of individuals of modern japonica. The lack of diversity in modern varieties is evidence of substantial genomic erosion, and it is likely to have negative coevolutionary consequences for the long-term sustainability of disease-resistant rice.

We show here that balancing selection is an important driver of the high diversity in NLRs of landraces. Both in modern varieties and landraces, balancing selection has elevated nucleotide diversity at the NLRs relative to that elsewhere in the genome. However, after controlling for differences in demographic histories, we found a greater number of genes under balancing selection across indica landraces than in the modern varieties. We also found that, compared to landraces, modern varieties are depleted in ancient polymorphisms that have been maintained by balancing selection. This observation is consistent with the evidence of stronger balancing selection (i.e., higher the selection coefficient γ), and with the fact that several NLRs were only found to be under balancing selection in indica landraces.

In addition to genes under balancing selection (i.e. genes with an excess of nucleotide diversity) being exclusively found in indica landraces, we also identified three genes under recurrent directional selection (i.e. genes with an excess of non-synonymous substitutions) that were unique to this group. The number of genes under strong directional selection is likely underestimated as only 285 NLRs displayed outgroup data and could thus be included in the analysis. Regardless of this limitation, we were surprised to find fewer signals of directional than balancing selection in NLRs in general, as modelling of finite populations also suggests that signatures of directional selection are more likely to be observable than signatures of balancing selection^33^. We also did not expect to detect evidence for directional selection only in landraces and not in modern varieties. Naively, one might expect arms race coevolution to be more widespread in modern agroecosystems, where NLRs with new resistance specificities are deployed and quickly overcome (boom-and-bust dynamics).

However, such coevolution could also play a role in traditional agroecosystems, where farmers have used the same landraces for centuries, being able to select from a multitude of landraces. The resistance breaking of a single NLR allele is then of little concern because there are other varieties with different alleles that confer resistance. The susceptible allele may then be replaced, or at least reduced in prevalence, leaving a signature of directional selection, but without risk to rice harvest. Although this is a plausible hypothesis, we must note that these differences in numbers of NLRs under positive selection could partly be because our ability to detect directional selection may be reduced in modern varieties due to their lower diversity.

An interesting case of an NLR under balancing selection in the landraces is RGA4. RGA4 is a helper NLR that interacts functionally with the sensor NLR RGA5, which recognizes the *Magnaporthe oryzae* effectors AVR-Pia and AVR1-CO39 or with the sensor NLR Pias-2 that detects the sequence-unrelated *M. oryzae* effector AVR-Pias^34-37^. RGA5 carries a non-canonical heavy metal-associated (HMA) domain after its LRR that directly binds AVR-Pia and AVR1-CO39 which is crucial for their detection^38,39^. Pias-2 harbors a completely different integrated domain^40^ of unknown function (DUF761) whose role in effector recognition remains unknown^37^. The canonical NLR domains of RGA5 and Pias-2 have limited sequence similarity (59.8 % identity) while the *RGA4* alleles coupled either with *RGA5* or with *Pias2* are highly identical (96.6 %) and functionally interchangeable. The *RGA4*/*RGA5* and *Pias1*/*Pias2* haplotypes also occur in wild rice species together with four additional *RGA5* alleles that have even other integrated domains^37^. Population genetics and comparative genomics analyses indicate that balancing selection maintains these multiple *RGA4/RGA5* alleles with contrasting recognition specificities across speciation in multiple species of the *Oryza* genus^35,37^. Our study shows that such balancing selection occurs also at the population level, within landraces, thereby potentially providing complementary protection against isolates with different virulence effectors. Interestingly, previous work has shown that in the Yuanyang terraces the effector AVR-Pia is absent from most *M. oryzae* isolates collected on indica landraces while it occurs at high frequency in isolates from japonica rice, on which it confers a gain in virulence^9^. Here, we report that RGA4 is present at high frequency and under strong balancing selection in indica landraces, with high diversity in the LRR domain. Under the interaction model described above, it is probably RGA5 or Pias-2 that are the main targets of coevolutionary interactions with fungal effectors and the signatures of balancing selection detected in RGA4 are a byproduct that results from compensatory changes in the helper induced by coevolution-driven changes in the sensors. The lack of variation detected in the CC domain is consistent with the fact that RGA4 and RGA5 interact through this domain^36^. Why then would compensatory changes take place in the LRR? One possibility is that RGA4 is not solely a helper NLR and has some other important function related to the response to other effectors, and thus that the observed changes in LRR are in fact not all driven by selection on its head-to-head partner.

Rice NLRs show not just SNP variation, but also presence/absence variation. Our data are summarized in Fig 3. When all accessions are compared, a slight majority of NLRs (∼ 350 / ∼ 600) are present in all investigated accessions (Fig 3A, Fig 3C-note the log scale in Fig 3C). A few NLRs are present in a minority of accessions. Other researchers have classified the conserved and presence/absence variable NLRs as “core” and “dispensable” ^41^, though we would argue that since polymorphism for recognition capacity underpins resistance and selects against specialist races of *Magnaporthe oryzae*^*42,43*^, the term “dispensable” conveys a misleading impression of lack of utility. Remarkably, presence/absence variation within land races mimics SNP variation, in that variation in number of NLRs within land races was higher than in japonica or indica varieties.

In aggregate, our work shows that rice NLRs represent a highly variable gene family, and that this variability is particularly high in landraces from the Yuanyang terraces. We found hints in the data for positive selection, but indications of balancing selection were more evident and pervasive than indications of directional selection. Therefore, the data tend to provide more support to the trench warfare hypothesis over the arms-race hypothesis as a general coevolutionary model for this class of genes. Integrated domains and LRRs seem to be the preferred target of balancing selection, consistent with their role in the recognition of pathogen-derived ligands. The effect of trench warfare is visible in the maintenance of high values of the π_max_ statistic in the landraces, which indicates the maintenance of ancient NLR alleles in these populations. Understanding how elevated NLR diversity and enrichment in older alleles reduce the burden of disease in traditional agroecosystems may help re-engineering modern crops and agroecosystems to make them less conducive to extant and emerging diseases.

## Materials & Methods

### Design of a rice RenSeq bait-library and application to explore NLR diversity

RenSeq analysis was performed on 49 rice accessions (Supplementary Table 1), representing seven indica landraces (36 accessions), two japonica landraces (two accessions) and eleven modern varieties of indica, japonica and aus. The nine landraces were represented by thirty-eight accessions, which are part of a rice diversity collection (single panicle descendants) established in 2014 and 2015 by sampling individual plants in the fields of traditional rice farmers in three villages (Xiaoshuijing, Xinjie, Qingkou) in the Yuanyang rice terrace region in Yunnan province (China). Thirty-one landrace accessions were selected as representatives of the genetic diversity of four major landraces cultivated in this region: Acuce (7 accessions), Baijiao (9 accessions), Hongjiao (8 accessions) and Xiaogu (8 accessions). Three accessions correspond to the Hongyang2 variety, a true breeding line bred from landrace germplasm and widely cultivated in the Yuanyang terraces in recent years. Three accessions are glutinous rice: Zinuo (indica), Huangpinuo (japonica) and Nuogu (japonica). Japonica rice is cultivated on limited surfaces in the Yuanyang terraces, ca. 5% of total rice acreage. The eleven modern varieties were selected in a collection^44^ representative of the world-wide rice phenotypic and genetic diversity (temperate japonica: four varieties, tropical japonica: three varieties, indica: four varieties, aus: one variety).

We designed a bait library capable of hybridizing to a wide variety of Asian rice NLRs. We characterized the NLR complements in the genomes of the japonica rice reference variety Nipponbare (MSU Rice Genome Annotation Project Release 7^45^) and the indica rice reference variety 93-11^46^ by three different approaches: (1) searching NBARC domain-coding sequences (containing Pfam|PF00931 motif) in the CDS of both genomes with PfamScan47; (2) identifying NLRs among CDS of both genomes with NLR Parser v1 ^48^using default parameters followed by removing those with an NBARC domain coding sequence shorter than 500 or longer than 1100 nucleotides; (3) recovering the NLR repertoires identified by Luo et al.^49^ in the Nipponbare and 93-11 genomes and filtering them for presence the NBARC domain (Pfam|PF00931). Redundancy within the japonica and the indica NLR gene sets was removed by positional information of the corresponding genes. In addition, to further remove redundancy in the NLR repertoire, NLRs whose NBARC -coding sequences were more than 95% identical between japonica and indica NLRs or among indica NLRs were removed by keeping the homolog with the longest NBARC domain. From the resulting set of 761 NLR sequences, 21,000 baits of 120 nucleotides and with 20 bp overlap were designed using a proprietary script from Arbor Bioscience (https://arborbiosci.com/). These oligos were aligned to the Nipponbare and 93-11 genomes with Blastn and oligonucleotides with more than 10 perfect matches per genome were excluded.

Genomic DNA was extracted from two weeks-old rice seedling using a CTAB method^50^. Enrichment and library preparation were carried out as described in Witek et al.^18,51^. Post-enrichment samples were sequenced using Illumina HiSeq 2500. We mapped RenSeq reads against a reference set of NLR sequences identified in Ensembl Plants Genes database (O. sativa indica ASM465v1 version 43, O. sativa japonica ASM465v1 version 45) using the BioMart utility to filter gene IDs with Interpro entry IPR002182 (NBARC domain). To avoid redundancy among sequences caused by orthology between *O. sativa* indica and *O. sativa* japonica, and because 36/38 of the landraces included in our dataset are of indica type, we determined orthology relationships between *O. sativa* indica and *O. sativa* japonica sequences, and retained a final set combining all *O. sativa* indica sequences with *O. sativa* japonica sequences having no ortholog in *O. sativa* indica (Supplementary Table 8). Blast-n analysis revealed that 593 of the 596 reference sequences had a minimum identity of 69% with sequences used as baits (Supplementary Table 8).

Raw Illumina paired-end reads from 49 rice accessions were aligned to the FASTA file of 596 NLR gene sequences using bowtie v2.3.5^52^ with the option –very-sensitive and the rest all defaults to produce 49 BAM files that were then sorted using Samtools (v1.9) tool^53,54^.

Illumina sequencing of captured RenSeq sequences provided 3,702,690 pairs of paired-end reads per accession on average (standard deviation: 4,462,281; min: 937,145; max: 27,528;626; Supplementary Table 9; Supplementary Figure 12). Variability in the number of reads, in particular between landraces and domesticates, had an impact on the proportion of sites covered and the coverage (Supplementary Figure 13; Supplementary Figure 14). In order to reduce the impact of the heterogeneity in the number of reads in our analysis of presence/absence and nucleotide diversity, we standardized our dataset to the same level of average sequencing depth, by randomly subsampling 937,145 pairs of reads for landraces and 2,342,863 pairs of reads for domesticates (937,145 is the number of pairs of reads of the less deeply sequenced accession V11; 2,342,863 is 937,145*2.5). This procedure reduced coupling between the number of reads and sequencing depth statistics (Supplementary Figure 13; Supplementary Figure 14), as observed by computing the standard deviation of sequencing depth across NLRs, which decreased from 219.3 to 10.8 (Supplementary Figure 15 Supplementary Table 9).

NLRs present copy number variation, so a substantial fraction of heterozygous calls is expected to result from hidden paralogy. Alleles at the same NLR locus can also vary in their affinity to sequencing baits, which can also influence the detection of heterozygosity. To identify and remove erroneous calls caused by hidden paralogy while controlling for allele imbalance, we used a SNP caller that explicitly models these two features. SNP calling was carried out using Reads2snp 2.0^55-57^ using 2592 combinations (Supplementary Table 10) of the following parameters: min (minimal number of reads required to call a genotype), th1 (minimal posterior probability required to call a genotype), par (filtering for SNPs caused by hidden paralogy), th2 (maximal p-value required to reject a paralogous SNP), aeb (accounting for allelic expression bias), fis (inbreeding coefficient), bqt (filtering out positions of quality below threshold), rqt (filtering out reads of mapping quality below threshold). To select the best combination of SNP calling parameters, we computed the number of segregating sites (S) and inbreeding coefficient (Fis) at NLRs for the group of four indica varieties in each of the 2592 SNP sets and compared with the values obtained for six random samples of four indica accessions from the three thousand rice genome dataset, referred to as 3KRG indica reference datasets ^58^. We computed the Euclidian distance between (Fis,S) estimated for the indica group in the 2592 SNP sets and for the three thousand rice genomes (3KRG) dataset, and by minimizing this distance we selected the “best” (or most comparable) SNP dataset with the lowest deviation from the 3KRG dataset. SNPs occurring in NLRs in Ref. ^58^ were identified by mapping protein sequences of our reference set of NLR against their older version of the 93-11 genomic sequence using Exonerate^59^. Four hundred eighty-eight of our reference set of 519 NLRs could be identified. Summary statistics S and Fis were computed using the Python package scikit-allel v. 1.3.3^60^. On average, inbreeding coefficient in the 3KRG indica reference datasets was Fis=0.62 (standard deviation: 0.02) and number of segregating sites was S=14707 (standard deviation: 1059). The closest of the 2592 Reads2snp datasets was dataset 286 (Fis=0.63, S=15016), obtained with the following parameters: min=10, th1=0.95, par=1, th2=0.001, aeb=False, Fis=0.99, bqt=40, rqt=20 (Supplementary Figure 16.

Rice is a selfing species and accessions were subjected to single seed descent before sequencing, so the number of heterozygous calls per SNP was expected to be much lower than postulated by Hardy-Weinberg equilibrium. Although the -par option in Reads2snp removed most SNP calls caused by hidden paralogy, SNPs with excess heterozygosity within populations remained in the dataset. In particular, plotting observed heterozygosity (Hobs) against the minor allele frequency (p) revealed SNPs with no or very few homozygous alternate calls, distributed along the Hobs=2p line, likely caused by gene duplications present in certain accessions (Supplementary Figure 17). SNPs with excess heterozygosity were removed using the same criterion as in ref. ^58^, by filtering out, in each landrace and rice subspecies, sites where observed heterozygosity was more than ten times the most likely value for a given frequency and inbreeding rate (Supplementary Figure 17). After filtering, summary statistics computed across the six 3KRG indica reference datasets (average [standard deviation]: Fis=1 [0], S=10200 [898]) remained very close to those computed on Reads2snp dataset 286 (Fis=1, S=10051). Reference NLR sequence used for SNP calling represented 2,423,478 bp. After masking 56,894 paralogous calls and SNPs with excess heterozygosity, the selected SNP set included 41,422 SNPs, of which 40,530 were biallelic.

### Empirical distribution of genome-wide polymorphism

To generate a baseline against which to identify features of polymorphism in genes of interest, we used previously published data for 68 accessions previously characterized^61-63^ using genotyping-by-sequencing (GBS), representing nine landraces and 28 modern varieties (Supplementary Table 1). GBS reads were mapped using bowtie v2.3.5^52^ (option –very-sensitive) against genic sequences predicted in the reference *O. sativa indica* genome 93-11 (Ensembl Genomes 45). SNP calling was carried out using Reads2snp 2.0^55-57^ the same set of parameters as selected for RenSeq, but relaxing constraints on sequencing depth and mapping quality: min=3, th1=0.95, par=1, th2=0.001, aeb=False, Fis=0.99, bqt=10, rqt=10. Of the 99,576,191 bp of genic reference sequence, 30,918,075 bp were covered by at least three GBS reads passing quality filters. After masking 121,672 paralogous calls and SNPs with excess heterozygosity, the selected SNP set included 200,098 SNPs, of which 199,128 were biallelic. The number of non-NLR genes characterized was 25,102 on average, and ranged from 8,968 (temperate japonica) to 30,875 (Hongjiao).

### Presence absence variation

The depth at each position of a NLR gene in each rice cultivars was obtained from the sorted BAM files using command depth in samtools. This depth values at all positions in a NLR gene is used to calculate a mean depth across the NLR gene. This gave a mean depth for each NLR gene in each rice cultivars. Each NLR mean depth was then normalized by taking the overall mean from all NLR gene mean depths in each rice cultivar, divide each NLR gene mean depth by overall mean depth and then multiplied by 100. The formula used was: = (NLR mean depth/Overall Mean) x 100. Jack-knife estimates of the coefficient of variation were obtained using the astropy.stats package in Python.

### Population subdivision

Clustering and phylogenetic network analyses were performed on biallelic SNPs. Clustering in K ancestral populations was performed using sNMF^20^. The K value ranged from 2 to 15 and each sNMF run was repeated 10 times. Clumpak64 was used to process sNMF output. Neighbor-net networks were built using Splitstree 5^21^. Phylogeny of RGA4 using RAxML v. 8.2.12^65^.

### Polymorphism and divergence

Summary statistics of variation were computed using Python package Egglib 3 (https://www.egglib.org/) after generating pseudo-alignments using the table of SNPs (in VCF format) and reference sequences.

Fisher exact, Kruskal-Wallis and Mann-Whitney tests were computed with scikit_posthocs 0.6.6 and scipy 1.8.0 in Python 3.6. To estimate summary statistics at functional domains, the coordinates of functional domains was obtained using InterPro, as implemented in Ensembl’s Biomart.

### Assessing the impact of sampling effort on measures of molecular variability

To assess the capacity of RenSeq to measure genetic diversity at NLRs, we compared read mapping statistics and measures of sequence variability estimated from RenSeq with estimates obtained from GBS, using a rarefaction approach to overcome potential biases related to differences in sample size. Average nucleotide diversity reached 90% of its maximum value with a pseudo-sample size of with 23 randomly selected accessions (Supplementary Figure 18), indicating that the majority of nucleotide diversity at NLRs has been uncovered with RenSeq data. Haplotype richness, in contrast, reached 90% of its maximum value with 39 accessions (Supplementary Figure 18), suggesting that the molecular diversification of NLRs occurs not only by mutation but also by recombination and gene conversion^66^. Rarefaction analysis of GBS data revealed that our dataset is sufficient to reliably characterize genomewide levels of polymorphism (Supplementary Figure 18).

### Demographic modeling

To determine if patterns of variation at NLRs departed from selective neutrality, we performed coalescent simulations to correct for deviation from demographic equilibrium (i.e. constant population size). We used an approximate Bayesian computation (ABC) framework ^22^ to identify the historical demographic model accounting for most features of the data at GBS loci without invoking selection. The most supported model served as a null hypothesis to test for selective neutrality at NLRs. ABC relies on the comparison between summary statistics calculated from observed data and the same statistics calculated from coalescent simulations under different demographic models. We used the folded frequency spectrum as the summary statistics, as computed using the scikit-allel^60^. For each group of landrace and modern rice, five models were compared: (i) the standard neutral model determined by a single parameter N1, the effective population size, (ii) an instantaneous bottleneck model ^67^, parameterized by the initial effective population size N1, the start of the bottleneck T and the strength of the bottleneck ST, which corresponds to the time period during which coalescence events are collapsed, (iii) an exponential growth model parameterized by the initial effective population size N1, the final effective population size N2 (N2<N1), the start of population growth T1, the end of population growth T2 (T2>T1), and the growth rate being computed as log(N2 / N1) / (T2-T1), (iv) a two epochs population contraction model parameterized by the initial effective population size N1, the final effective population size N2 (N2>N1), the time of population change T. A two epochs population expansion model was initially included in analyses, but finally dropped as no simulations were accepted under this model for any dataset. Prior distributions are given in Supplementary Table 11. Simulations were performed using msprime68-70 assuming a recombination rate of 1e-8/generation/bp. Model selection and parameter estimation were performed using the R abc package ^71^. We simulated 1 million multilocus datasets for each model and population. Posterior probabilities of demographic scenarios were computed using the rejection (“rejection”) and the multinomial logistic regression method (“mnlogistic”) methods with a tolerance rate of 0.5%. Cross-validation for model selection based on 100 simulations per model at tolerance level 0.5% revealed that our ABC framework could efficiently distinguish between the different demographic models (Supplementary Table 12).

Goodness-of-fit analyses ^72^ indicated that models with the highest posterior probabilities provided a good fit to the data for all groups and populations (Supplementary Table 13. To further check the fit of the models with highest posterior probabilities to the observed, we performed posterior predictive checks ^71^. For the best supported model of each population/group, the posterior distribution of each parameter was binned in twenty classes, random values were sampled by randomly picking classes and drawing values assuming a uniform distribution within the class. One thousand datasets of the same sample size and sequence length as GBS sequences were simulated per population/group in msprime using the sampled multivariate parameters. For all populations/groups, the best supported models were able to reproduce the observed values of π and Tajima’s D, confirming their goodness-of-fit (Supplementary Figure 19).

Selective neutrality at NLR genes was tested by simulating null distributions using the most supported demographic models inferred from GBS data. To generate null distributions, 10000 datasets of the same sample size and sequence length as NLR sequences were simulated in msprime by sampling multivariate parameters from posterior distributions using the same procedure as for posterior predictive checks.

### Directional selection

The SnIPRE framework uses a generalized linear mixed model to estimate the influence of mutation rate, species divergence time, constraint, and selection effects on polymorphism and divergence. Genome wide effects are incorporated into the analysis as fixed effects, while individual gene effects are incorporated as random effects, which allows to combine information across genes and increases power to detect the effects of selection on a gene-by-gene basis. We focused our analyses on the selection effects, which reflect the selection coefficients (γ), and the constraint (or non-synonymous) effects, which reflects mutational constraint (1-f, f being the proportion of non-synonymous mutations that are not lethal).

## Data availability

RenSeq sequencing data are available in NCBI under accession number PRJEB23459. Single-nucleotide polymorphism datasets are available in Zenodo (doi: 10.5281/zenodo.7386472)

## Acknowledgments

We thank Peter Balint-Kurti for critical reading of the manuscript, and Dan MacLean for useful suggestions.

## Notes

https://zenodo.org/record/7386473#.Y424PezMJbY

https://www.ncbi.nlm.nih.gov/sra/?term=PRJEB23459

### Competing Interest Statement

The authors have declared no competing interest.

