## Supplementary figure 1 for "Extensive immune receptor repertoire diversity in disease-resistant rice landraces"

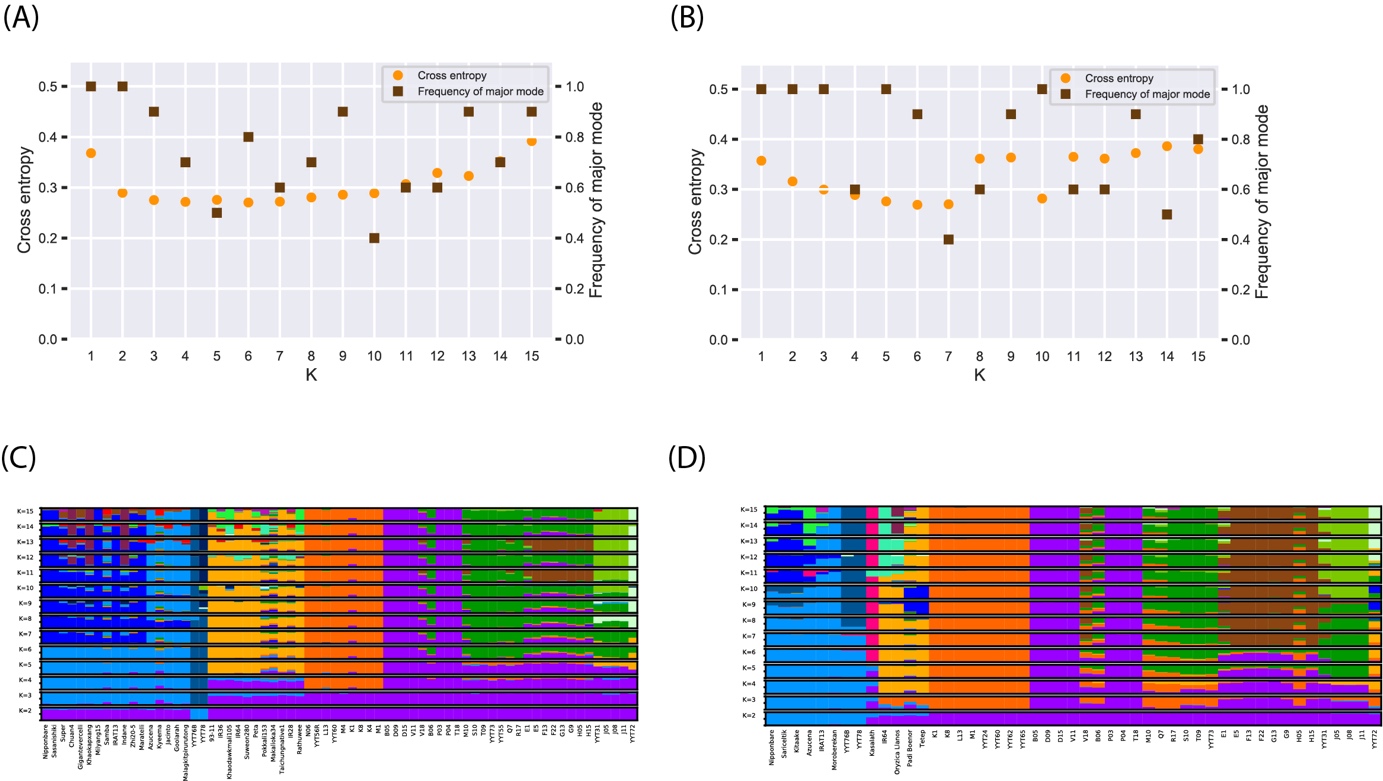

Supplementary Figure 1. Population subdivision was inferred with the sNMF program^1^ from 49 accessions, representing 13 varieties or landraces shown with different colors. (A) (B) Cross-entropy and frequency of the main clustering solution (i. e. major mode) as a function of the number of clusters K modeled in sNMF analyses of population subdivision for GBS and RenSeq data, respectively. (C) (D) Ancestry proportions in *K*=2 to K=15 clusters, for GBS and RenSeq data respectively, as estimated with sNMF software; each accession is represented by a vertical bar divided into *K* segments indicating membership in *K* clusters. For RenSeq data, analysis was based on 31,770 biallelic SNPs with less than 80% missing data. For GBS data, analysis was based on 60,166 biallelic SNPs with less than 50% missing data.

1	Frichot, E., Mathieu, F., Trouillon, T., Bouchard, G. & François, O. Fast and efficient estimation of individual ancestry coefficients. *Genetics* **196**, 973-983 (2014).
