## Supplementary figure 2 for "Extensive immune receptor repertoire diversity in disease-resistant rice landraces"

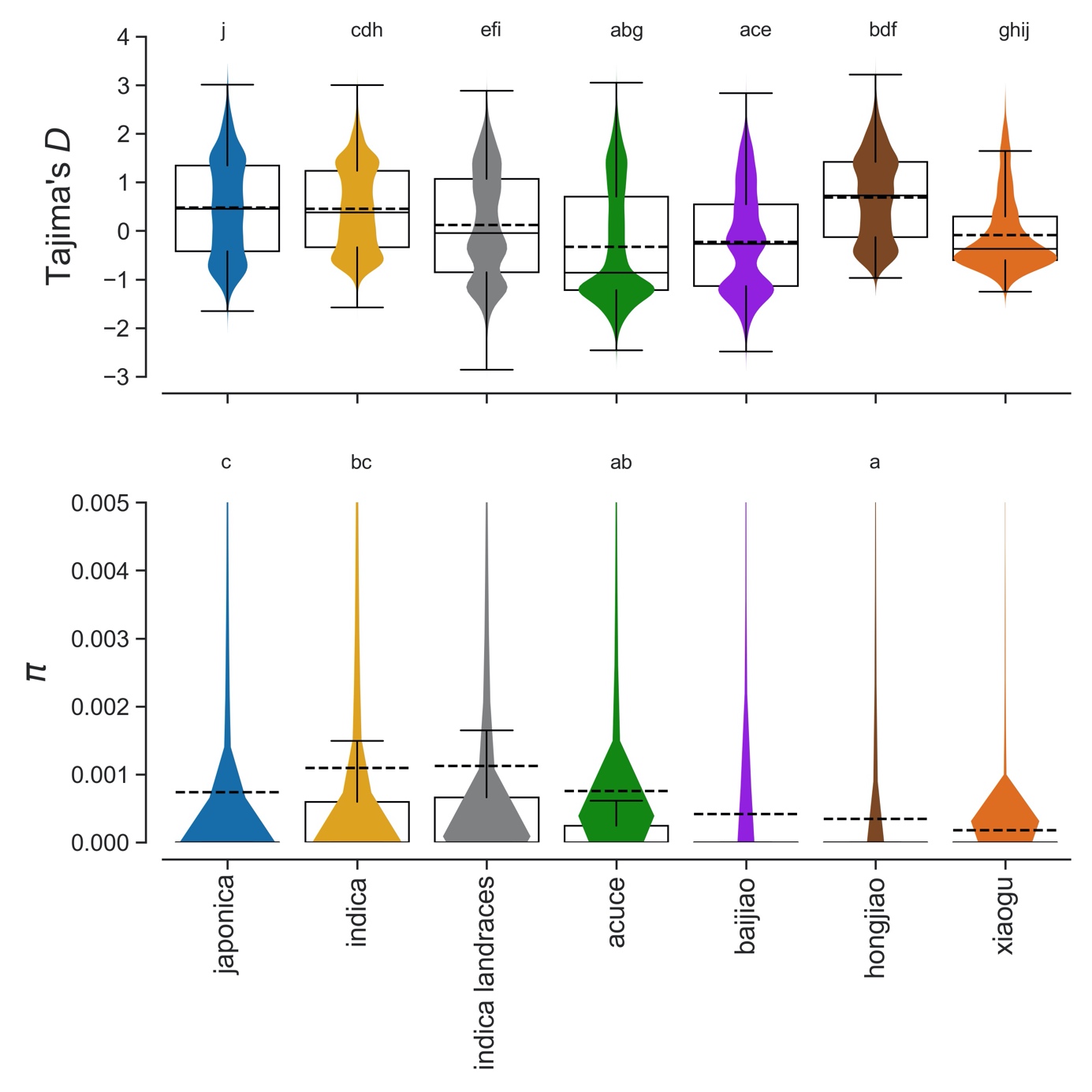

Supplementary Figure 2. Violin plots and box plots of nucleotide diversity (π) and Tajima’s D. The ‘indica landraces’ group includes one randomly chosen accession per individual landrace (29 other resamplings were carried out but results are not presented here as they are essentially the same, see Supplementary Table 1). Kruskal-Wallis tests were all statistically significant (Tajima’s D: H=3911.4, d.f.=6, p<0.0001; π: H=11330.1, d.f.=6, p<0.0001). Shared superscripts indicate non-significant differences (Mann-Whitney posthoc tests, p>0.05, Holm-Bonferroni correction). In box plots, dashed black line is mean, solid black line is median. The y-axis of the nucleotide diversity was cropped for visually optimal presentation, but all points were included in statistical tests.
