## Supplementary figure 3 for "Extensive immune receptor repertoire diversity in disease-resistant rice landraces"

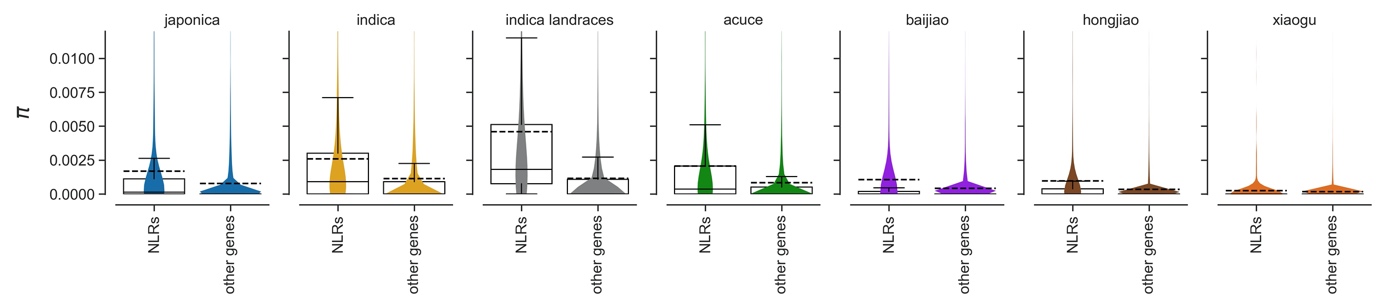
 Supplementary Figure 3. Box- and violin-plots of nucleotide diversity π in RenSeq data (NLRs) and GBS data (other genes). The ‘indica landraces’ group includes one randomly chosen accession per individual landrace (29 other resamplings were carried out but results are not presented here as they are essentially the same, see Supplementary Table 1). In box plots, dashed black line is mean, solid black line is median. The y-axis was cropped for visually optimal presentation.
