## Supplementary figure 4 for "Extensive immune receptor repertoire diversity in disease-resistant rice landraces"

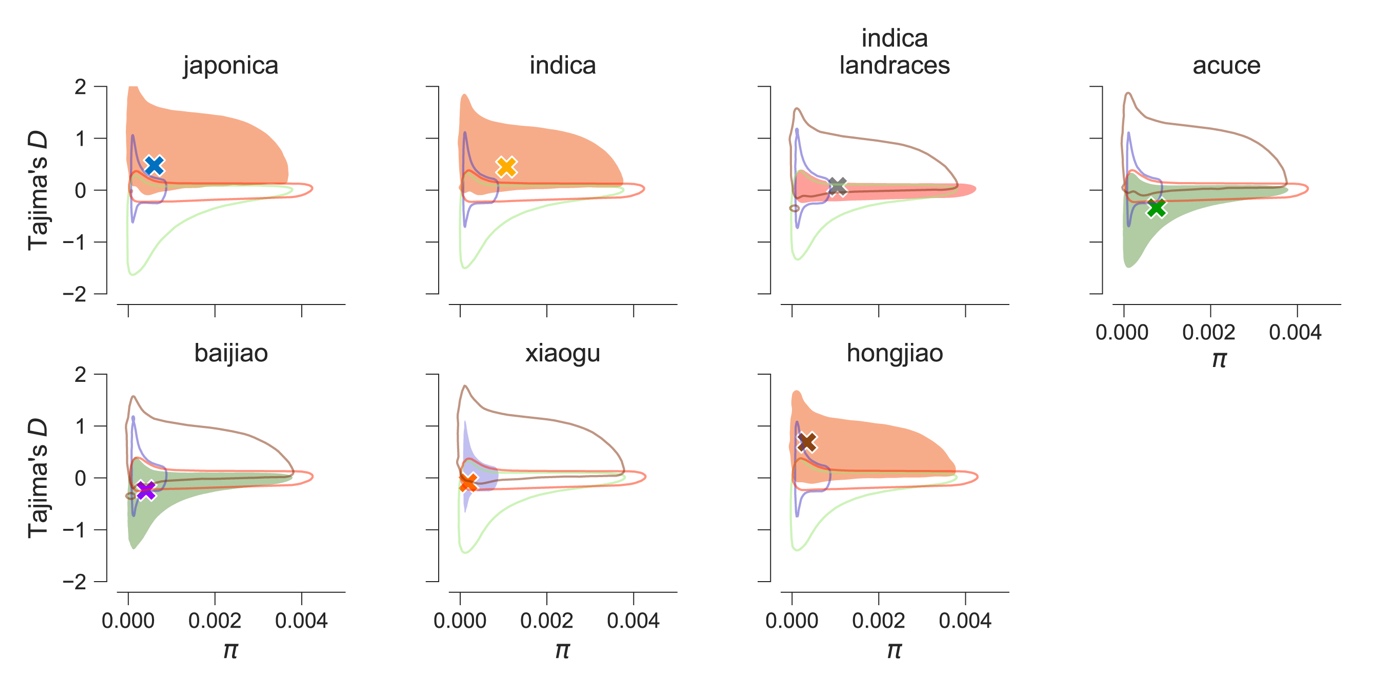

Supplementary Figure 4. Observed and simulated Tajima’s D and nucleotide diversity π in approximate Bayesian computations. Simulations were conducted under four different models. Crosses represent observed values of summary statistics. Summary statistics computed on one million simulated datasets are represented as kernel density estimate plots, as implemented in Python package Seaborn 0.11.2. The area between bivariate contours is filled in for models with highest posterior probabilities (Supplementary Table 3).
