## Supplementary figure 5 for "Extensive immune receptor repertoire diversity in disease-resistant rice landraces"

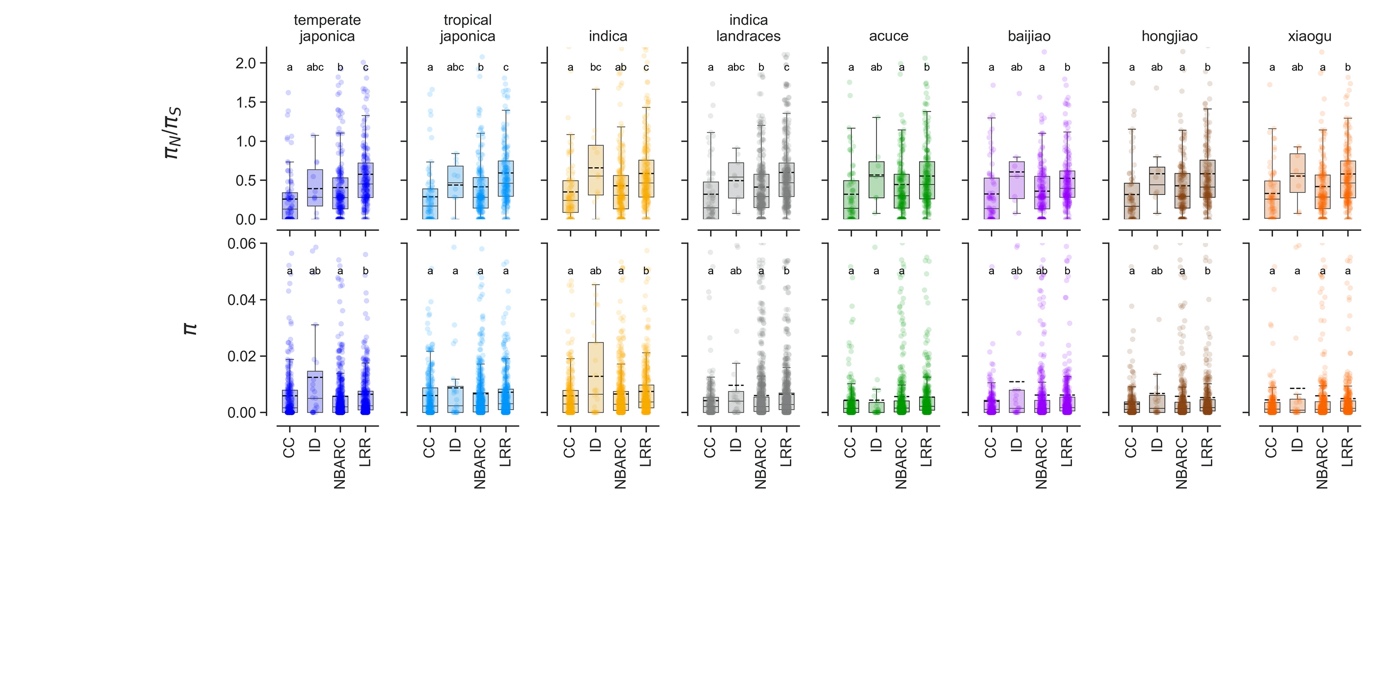

Supplementary Figure 5. Summary statistics per NLR in functional domains. Genes with all four domains without polymorphism were excluded. Shared superscripts indicate non-significant differences (Mann-Whitney posthoc tests, p>0.05, Holm-Bonferroni correction). A number of data points were cropped for visually optimal presentation, but included in statistical tests. In box plots, dashed black line is mean, solid black line is median.
