## Supplementary figure 6 for "Extensive immune receptor repertoire diversity in disease-resistant rice landraces"

(A)

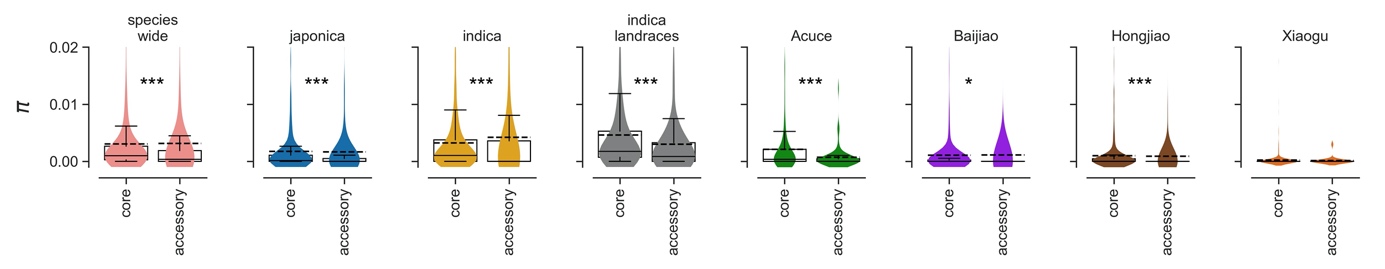

(B)

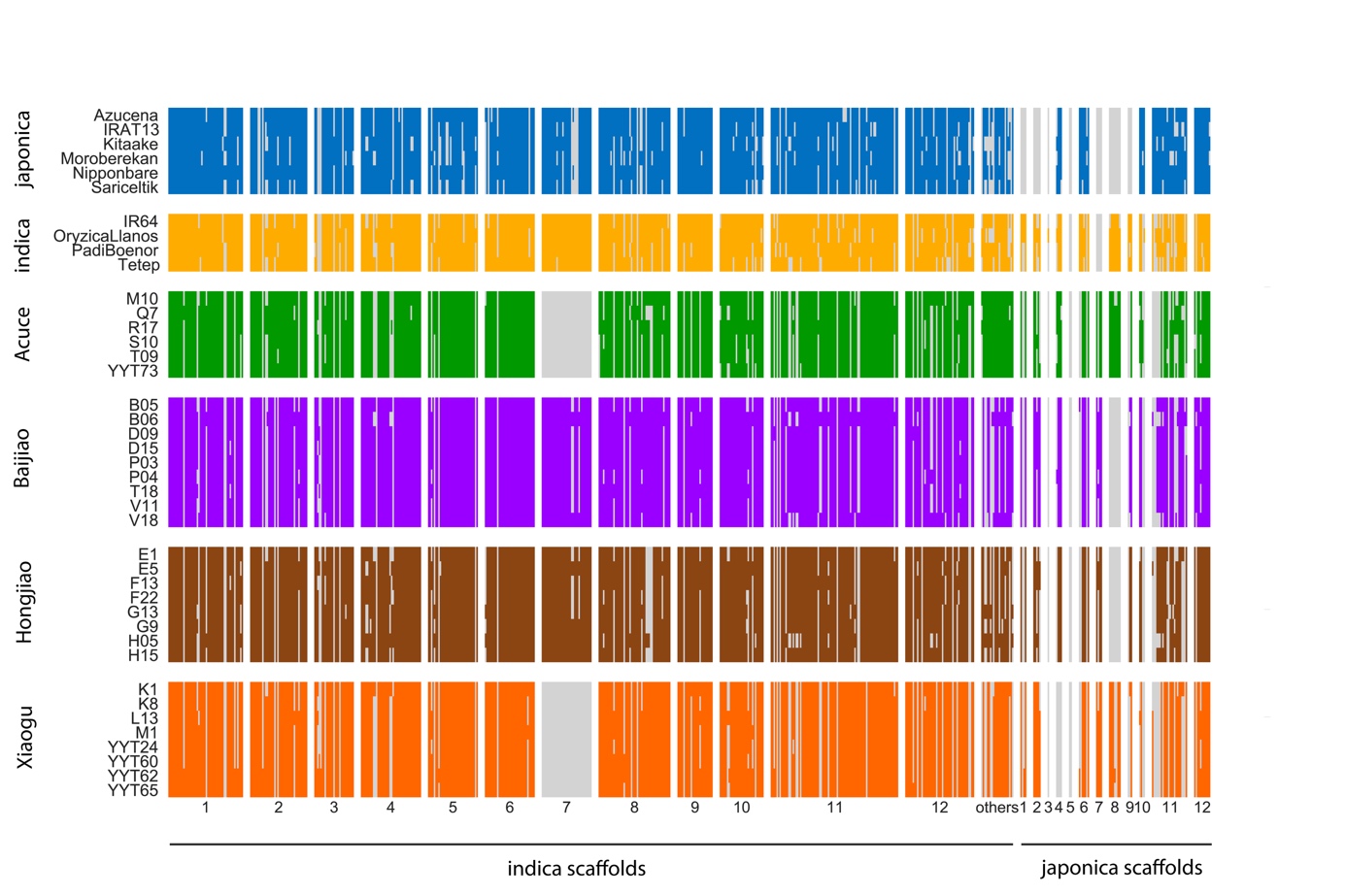

(C)

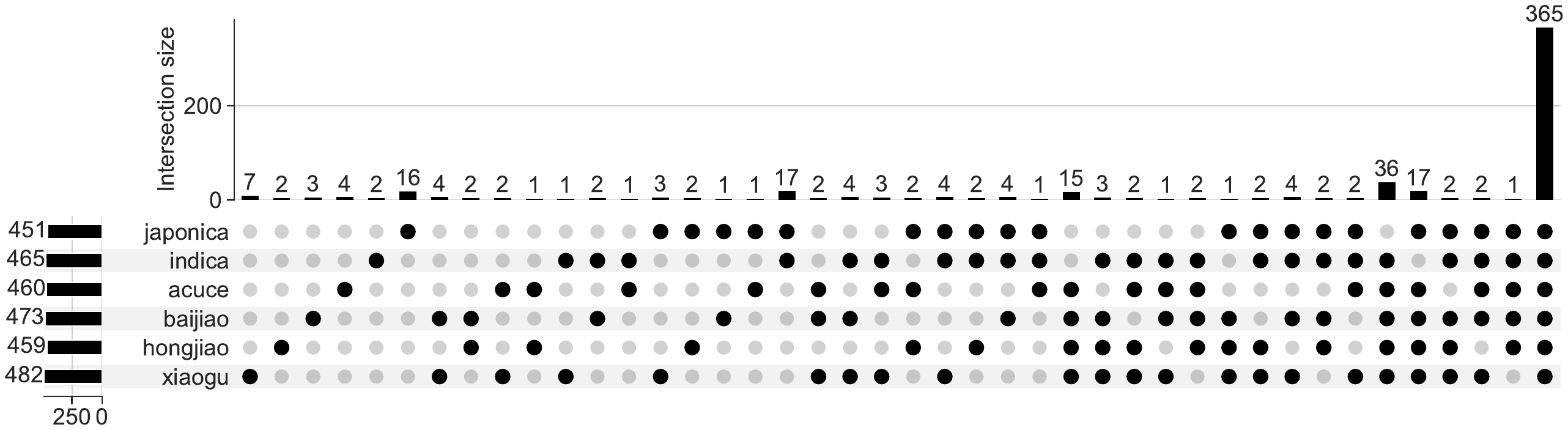

(D)

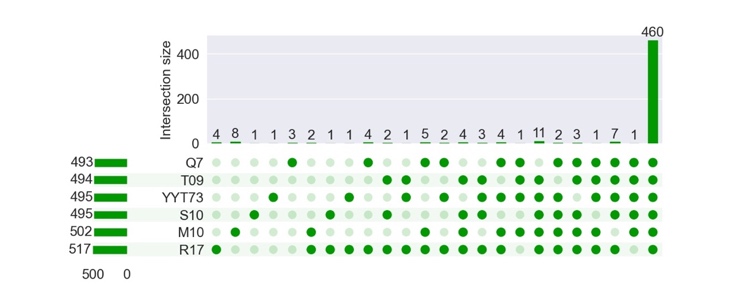

(E)

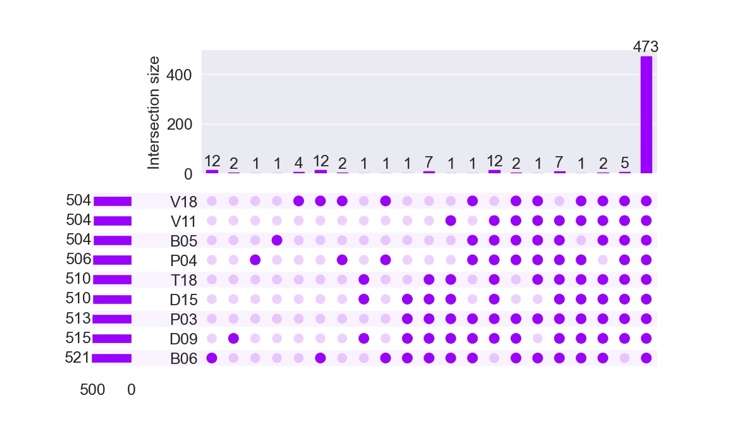

(F)

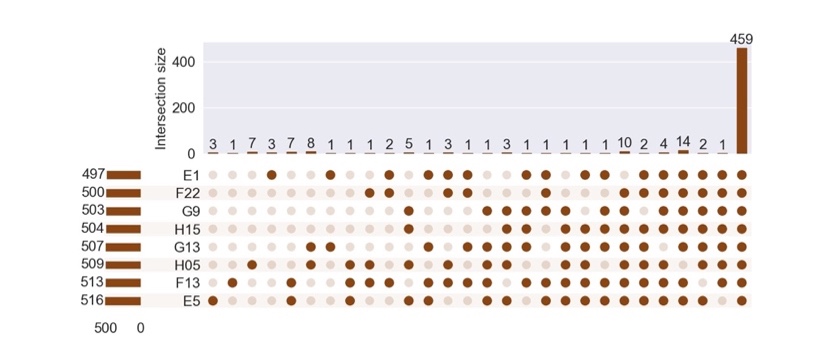

(G)

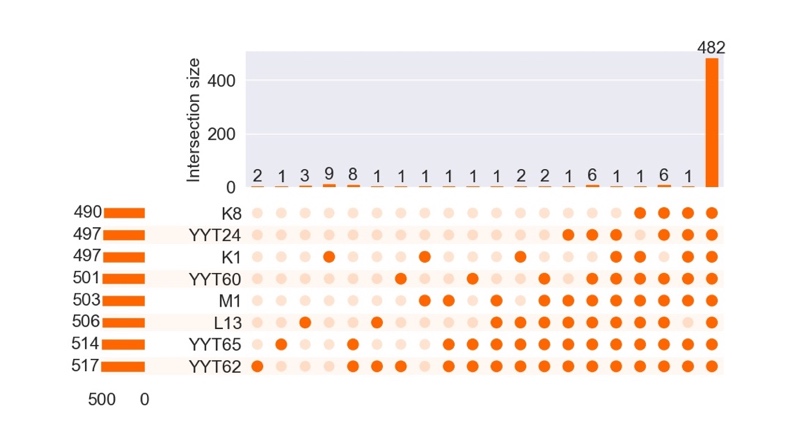

(H)

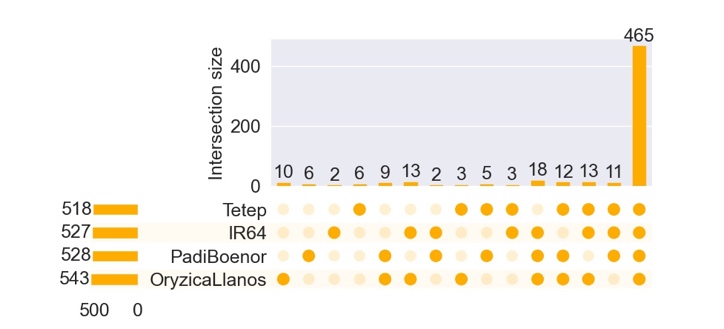

(I)

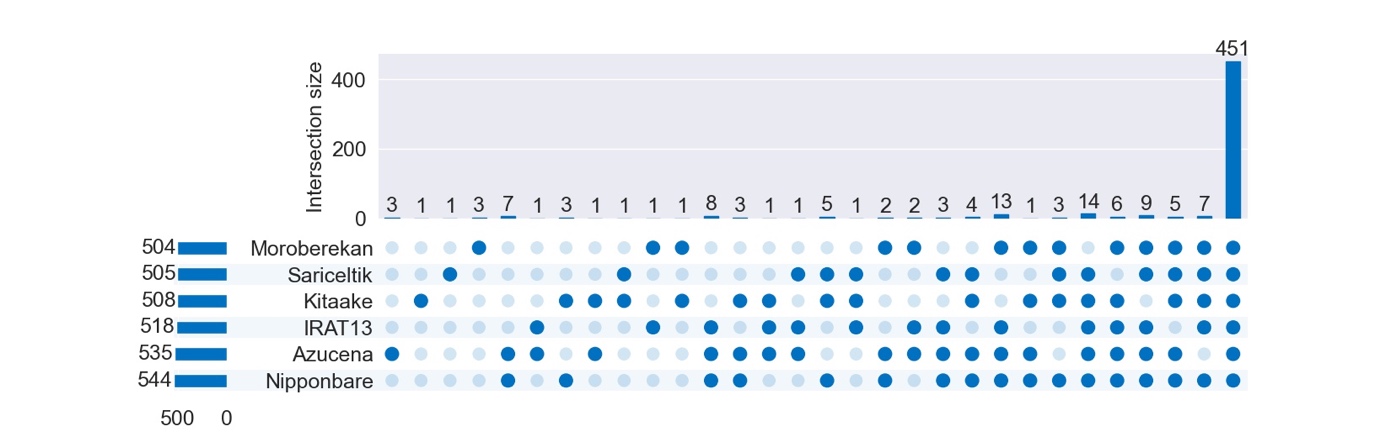

Supplementary Figure 6. (A) Nucleotide diversity π in core and accessory NLRs. ***p<0.001, *p<0.05, Mann-Whitney posthoc tests, Holm-Bonferroni correction. In box plots, dashed black line is mean, solid black line is median. (B) Distribution of NLRs in six rice populations, with absence in grey and presence in color; NLRs are organized according to their order in the indica and japonica reference assemblies (C). Distribution of core NLRs in the whole dataset. Vertical barplot represents the distribution of NLRs that belong to the core genome in at least one population. Horizontal barplot represents the number of core NLRs per population. A core is a NLR that is present in all accessions of all subsamples of two accessions from a given population. Presence of a core NLR is indicated by a black disk, absence by a grey disk. (D-I) Distribution of NLRs in populations. Vertical barplot represents the distribution of NLRs present in 1-N individuals, N. being the sample size. Horizontal barplot represents the number of NLRs per accession. Presence of a NLR is indicated by a coloured disk, absence by a smaller, shaded disk.
