## Supplementary figure 7 for "Extensive immune receptor repertoire diversity in disease-resistant rice landraces"

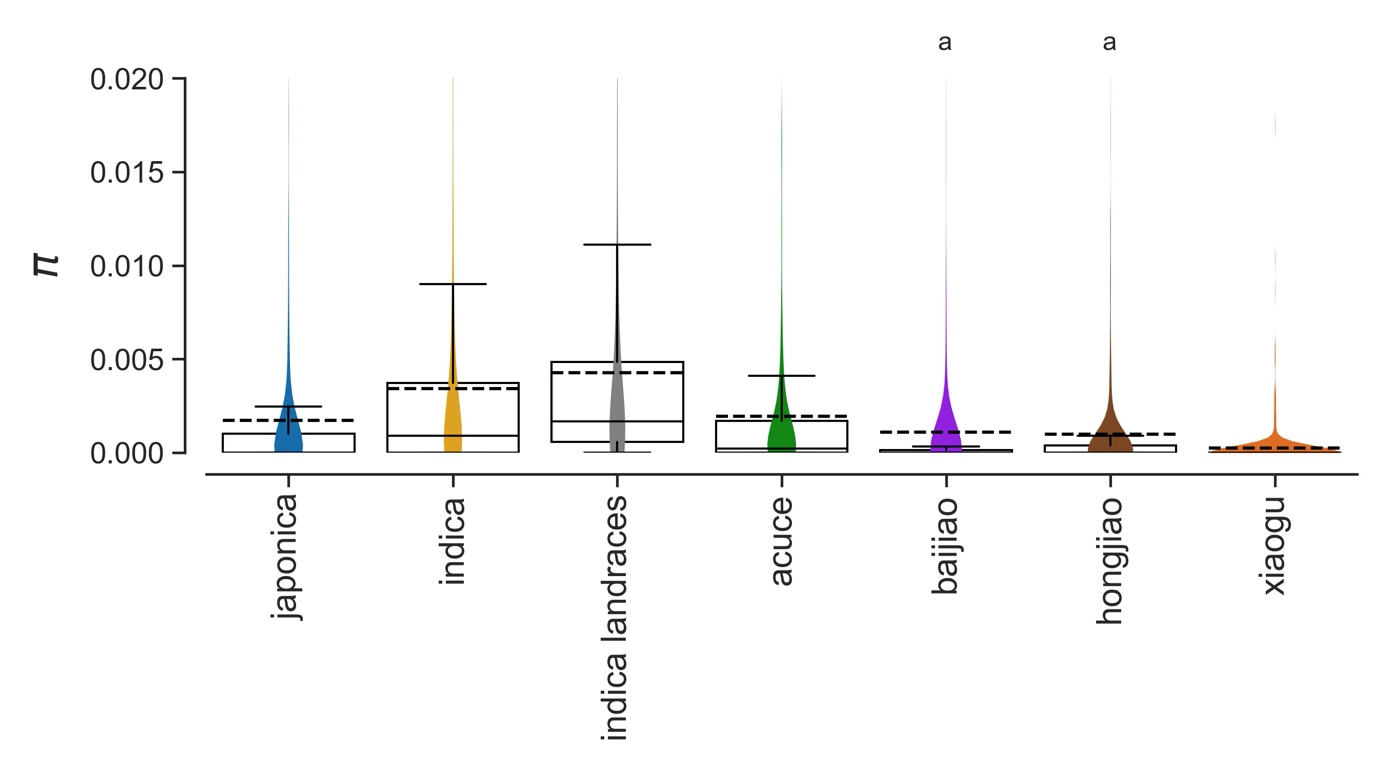

Supplementary Figure 7. Violin plots and box plots of nucleotide diversity (π) in NLRs characterized using RenSeq. The ‘indica landraces’ group includes one randomly chosen accession per individual landrace (29 other resamplings were carried out but results are not presented here as they are essentially the same, see Supplementary Table 1). Shared superscripts indicate non-significant differences (Mann-Whitney posthoc tests, p>0.05, Holm-Bonferroni correction). In box plots, dashed black line is mean, solid black line is median. The y-axis was cropped for visually optimal presentation, but all points were included in statistical tests.
