## Supplementary figure 8 for "Extensive immune receptor repertoire diversity in disease-resistant rice landraces"

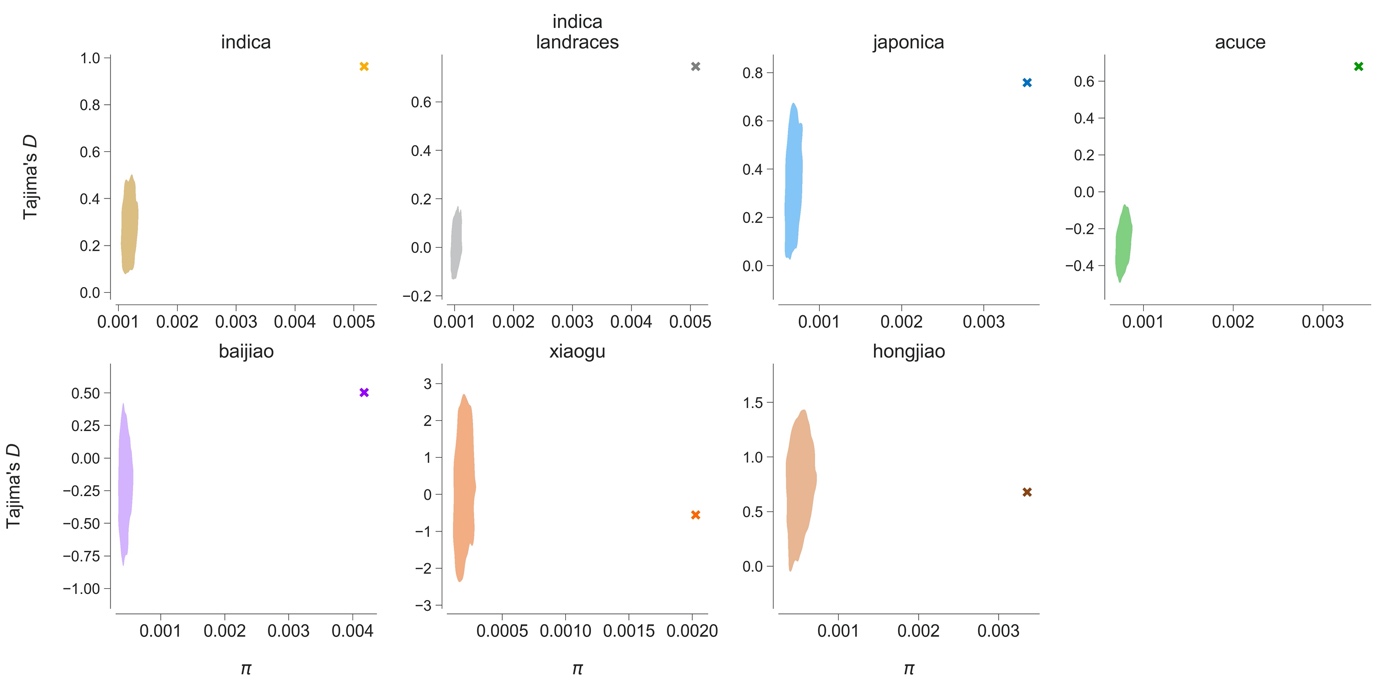

Supplementary Figure 8. Testing for selective neutrality at NLR genes using most supported demographic models inferred from GBS data to simulate null distributions. Crosses represent observed values of summary statistics Tajima’s D and nucleotide diversity π at NLRs. Summary statistics computed on 10000 simulated datasets are represented as kernel density estimate plots, as implemented in Python package Seaborn 0.11.2. Datasets of the same sample size and sequence length as NLR sequences were simulated by sampling multivariate parameters from posterior distributions (see details in Methods section).
