## Supplementary figure 9 for "Extensive immune receptor repertoire diversity in disease-resistant rice landraces"

(A)

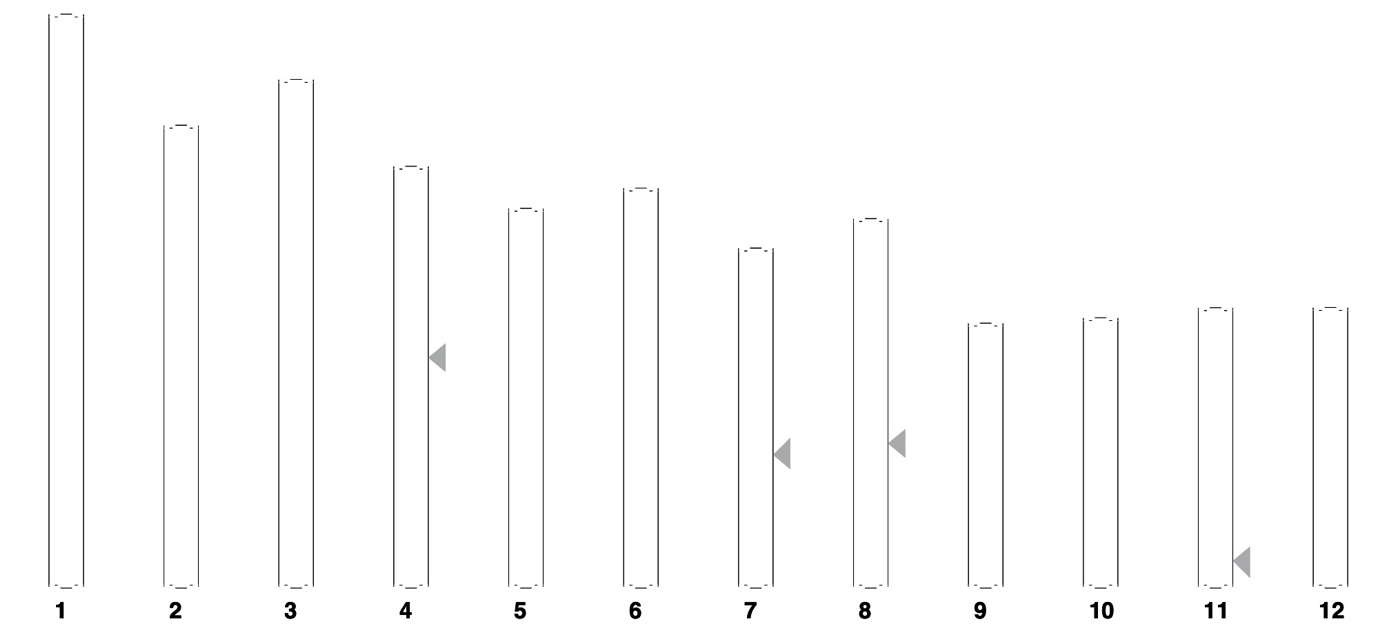

(B)

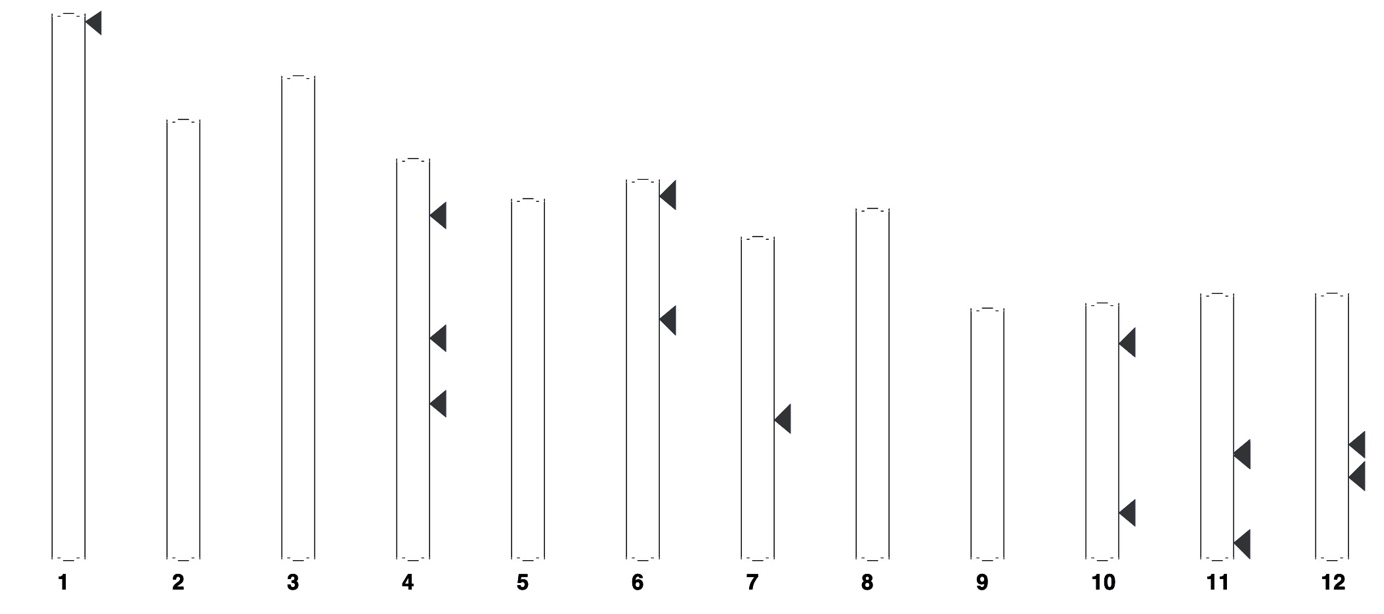

(C)

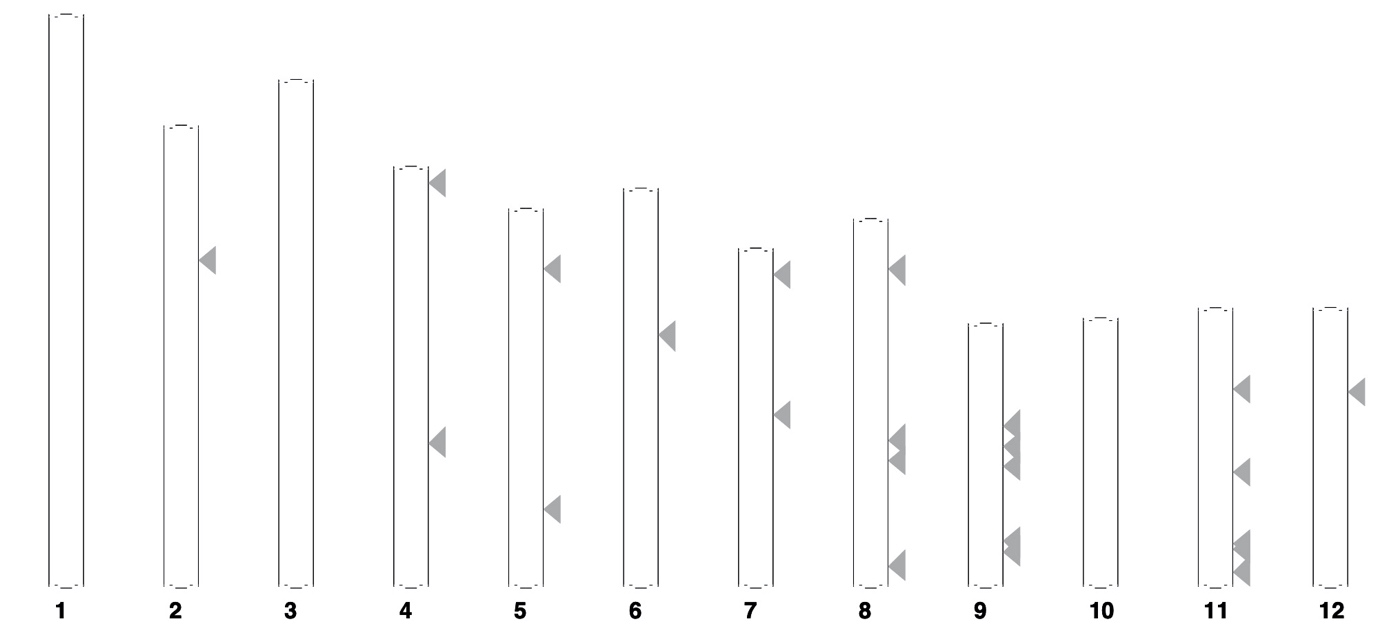

Supplementary Figure 9. Chromosomal location of NLRs under balancing selection in indica (A), japonica (B) and indica landraces (C) in the ASM465v1 indica reference genome.
