## Supplementary figure 10 for "Extensive immune receptor repertoire diversity in disease-resistant rice landraces"

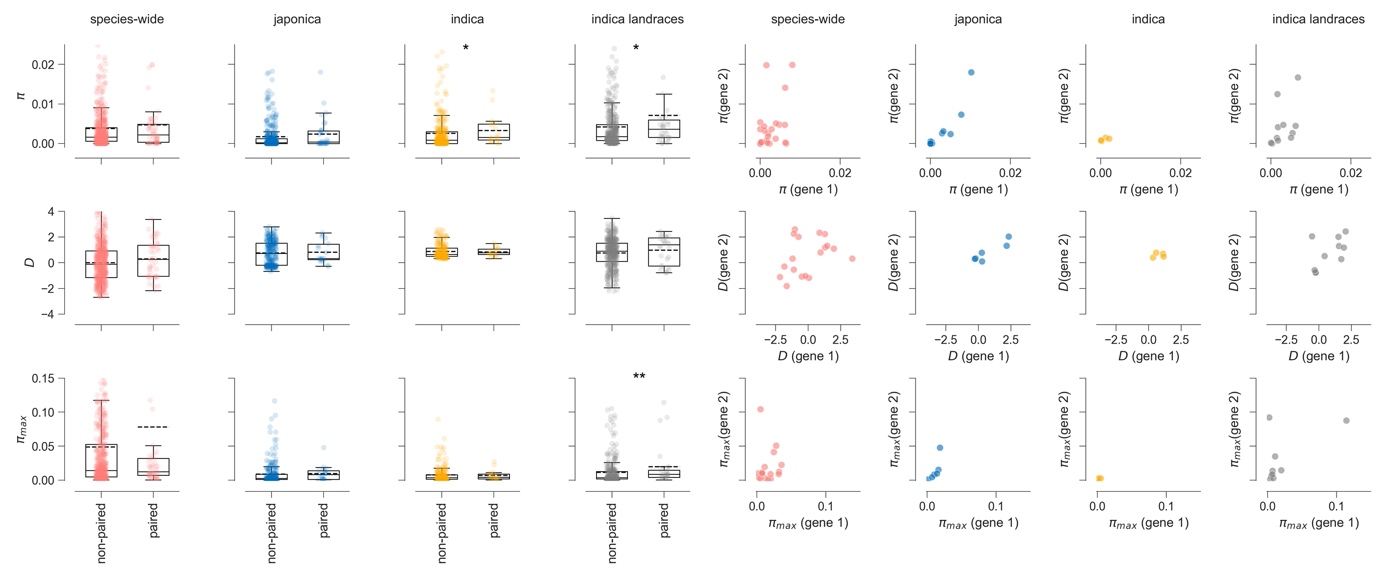

Supplementary Figure 10. Nucleotide diversity π, Tajima’s D, and the maximum number of pairwise differences π_max_ for non-paired NLRs and paired head-to-head NLRs. *p<0.05, **p<0.01, one-sided Mann-Whitney tests with Bonferroni-Holm correction.
