## Supplementary figure 11 for "Extensive immune receptor repertoire diversity in disease-resistant rice landraces"

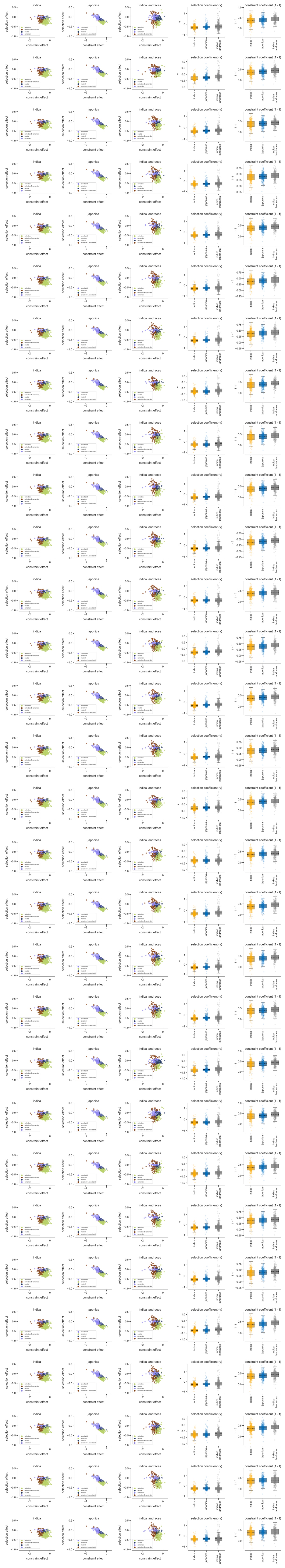

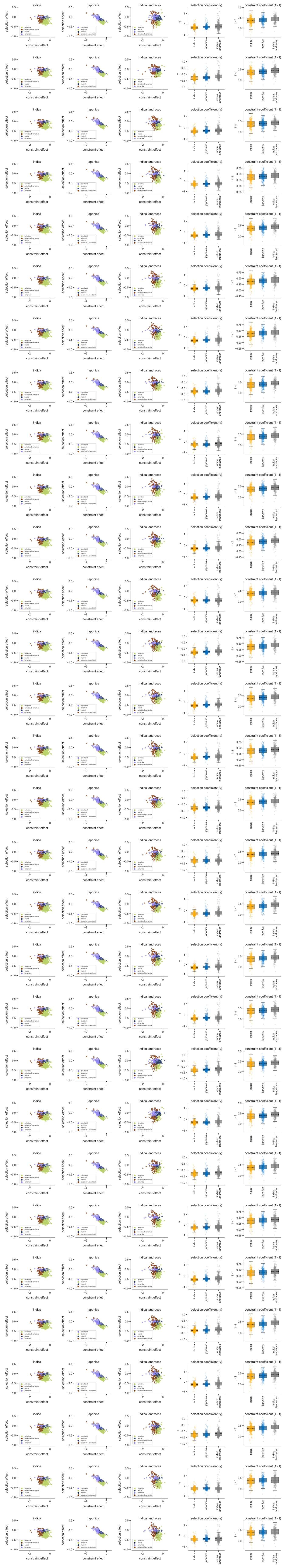

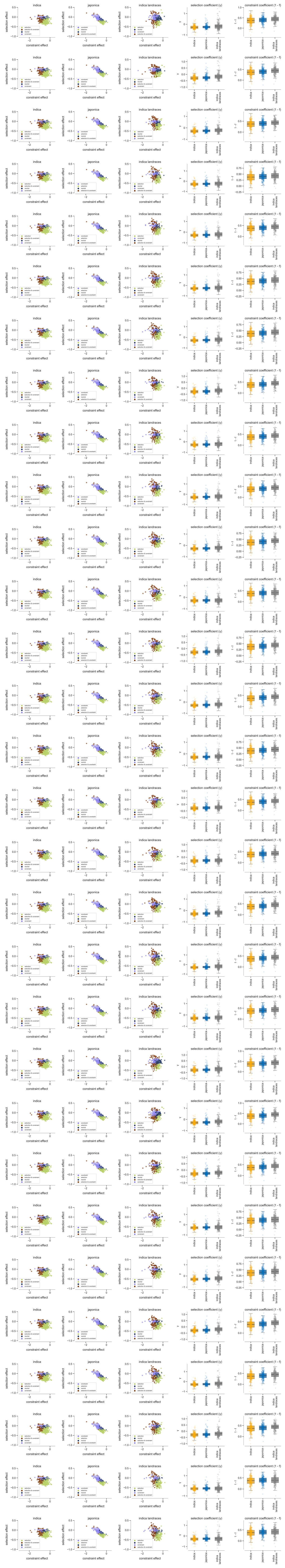

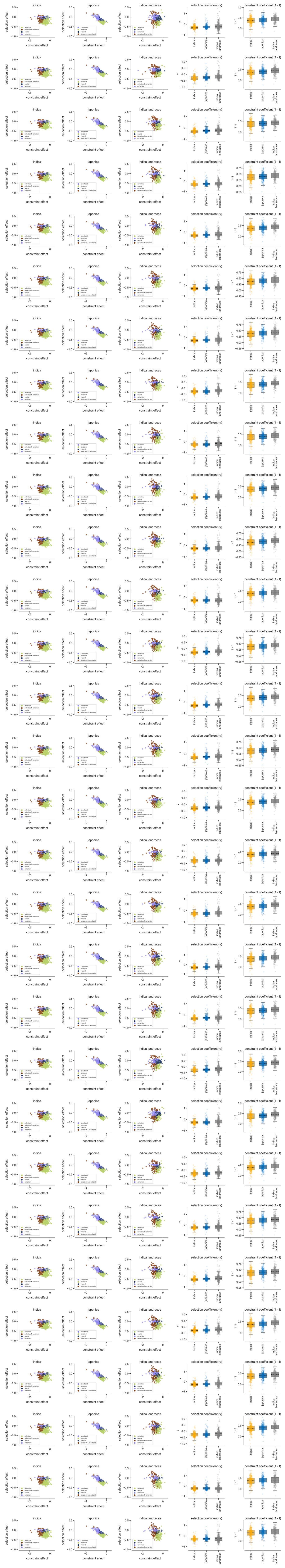

Supplementary Figure 11. SnIPRE estimates of recurrent directional selection in 285 polymorphicNLRs with outgroup data. One row per resample of 30 accessions, with one accession per landrace. For each row, panels represent: (i) Selection and constraint effects in indica, (ii) Selection and constraint effects in japonica, (iii) Selection and constraint effects in indica landraces, (iv) Selection coefficients in indica, japonica and indica landraces, (v) Constraint coefficients in indica, japonica and indica landraces. The selection effects reflect the selection coefficients (γ), with γ>0 indicating positive selection and γ<0 negative selection. The constraint (or non-synonymous) effects reflect mutational constraint (1-f, f being the proportion of non-synonymous mutations that are not lethal).
