## Supplementary figure 12 for "Extensive immune receptor repertoire diversity in disease-resistant rice landraces"

### Slide 1
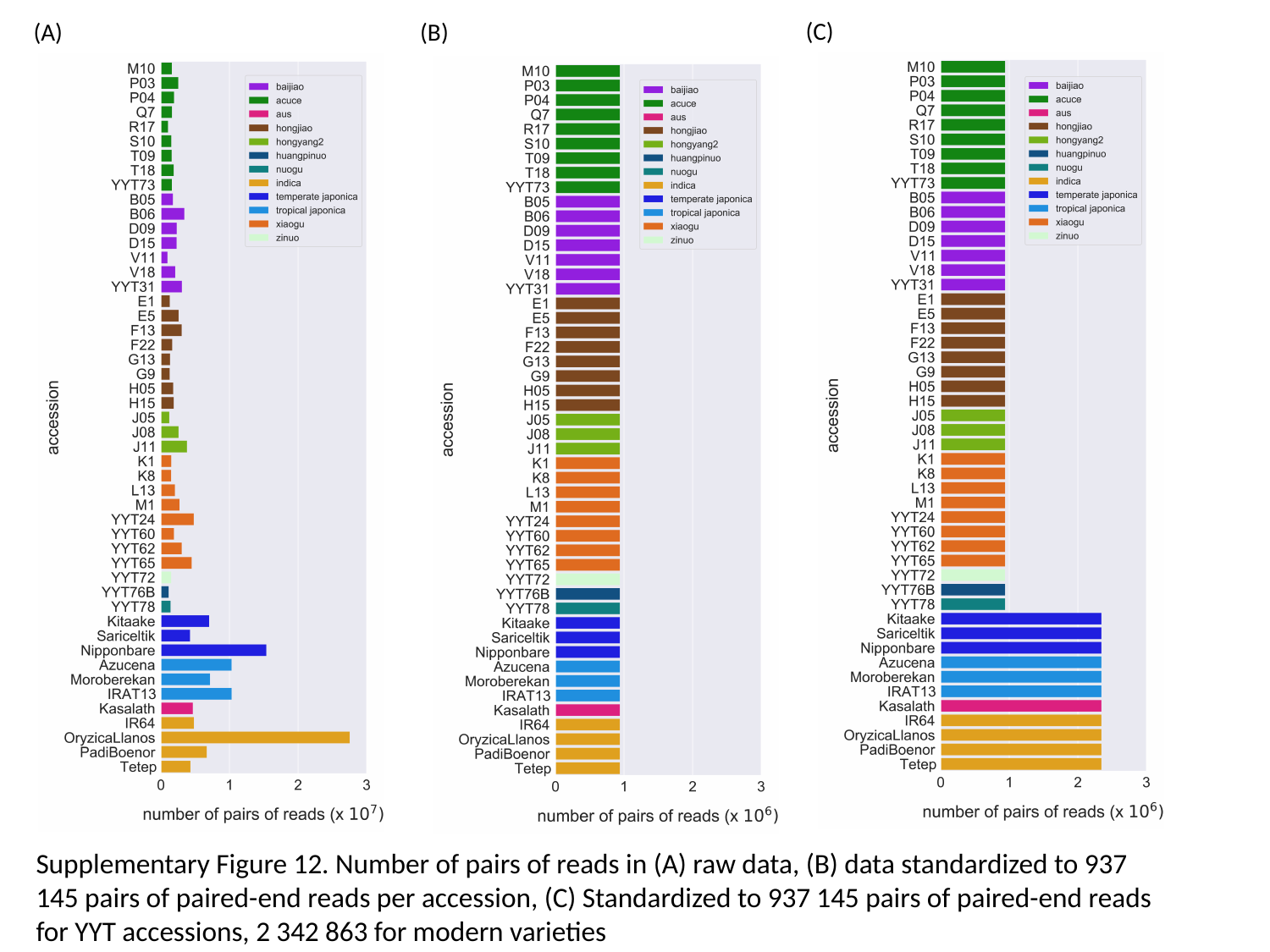

(C)
(A)
(B)
Supplementary Figure 12. Number of pairs of reads in (A) raw data, (B) data standardized to 937 145 pairs of paired-end reads per accession, (C) Standardized to 937 145 pairs of paired-end reads for YYT accessions, 2 342 863 for modern varieties
