## Supplementary figure 14 for "Extensive immune receptor repertoire diversity in disease-resistant rice landraces"

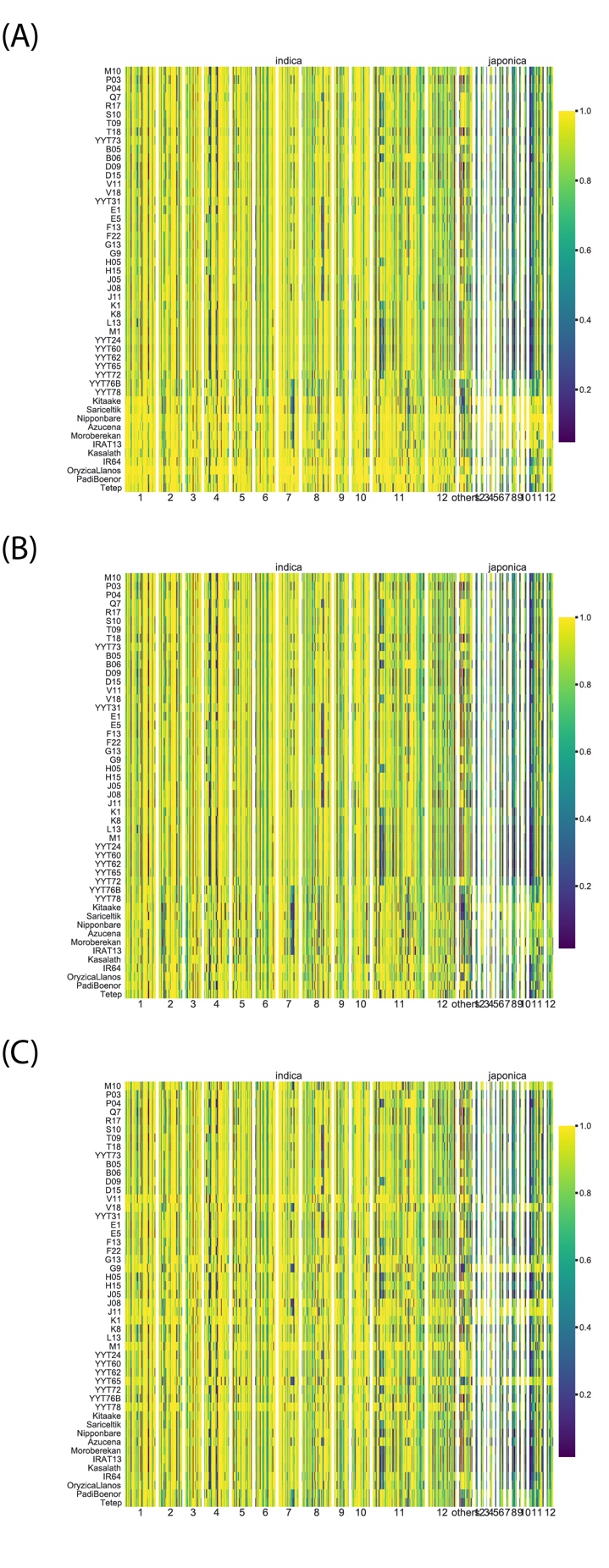

Supplementary Figure 14. Proportions of positions covered per NLR in (A) raw data, (B) data standardized to 937 145 pairs of paired-end reads per accession, (C) Standardized to 937 145 pairs of paired-end reads for YYT accessions, 2 342 863 for domesticates. NLRs appear in the same order as in the references Oryza indica Ensembl Genomes 43 and Oryza japonica Ensembl Genomes 45. NLRs are organized in 13 blocks in indica (12 main scaffolds, and pooling of smaller scaffolds), and 12 blocks in japonica (12 main scaffolds).
