## Supplementary figure 16 for "Extensive immune receptor repertoire diversity in disease-resistant rice landraces"

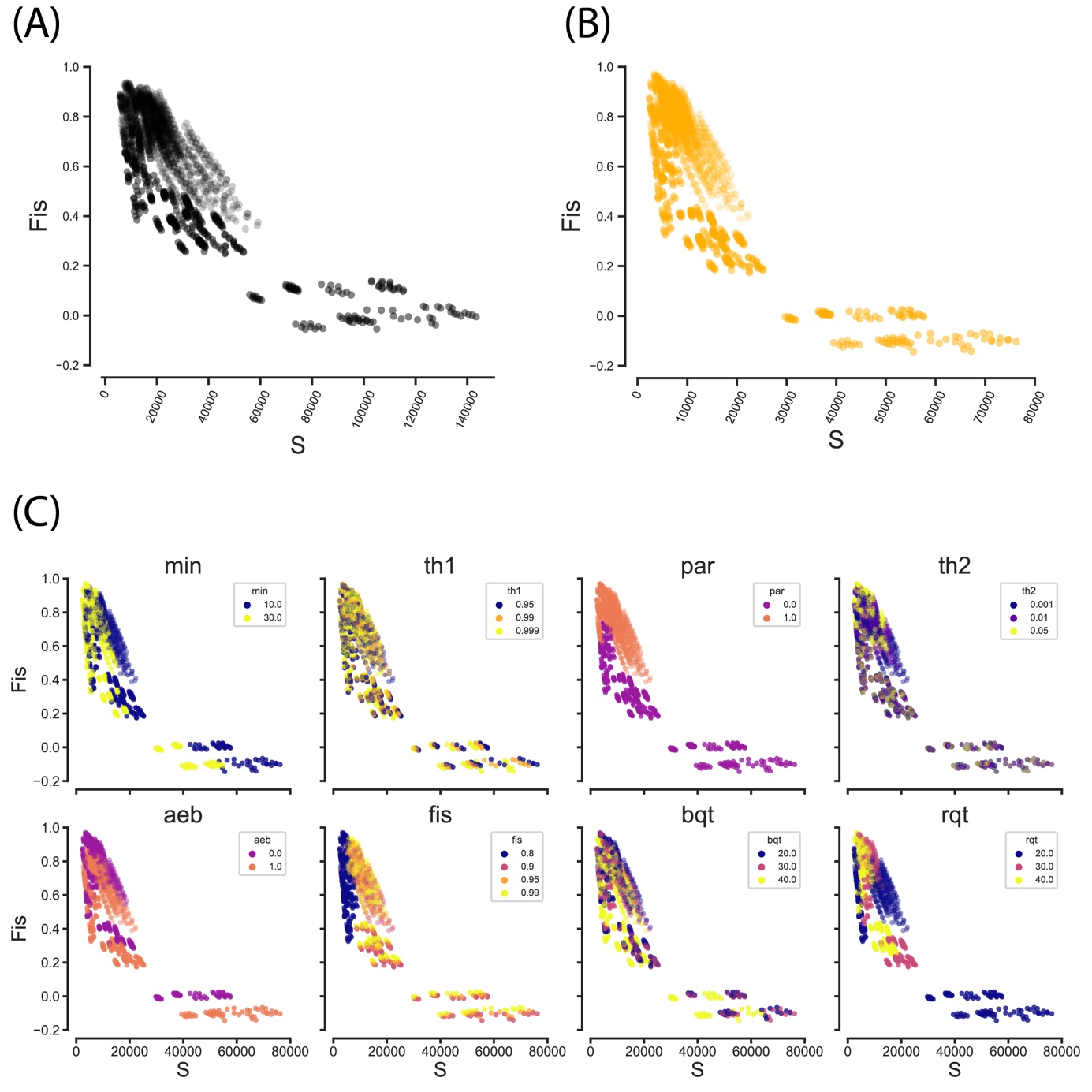

Supplementary figure 16. Inbreeding coefficient (Fis) versus number of segregating sites (S) in 2592 SNP sets generated with Reads2snp2. (A) Fis vs S calculated over the whole dataset, (B) Fis vs S for indica accessions, (C) Fis vs S calculated over the whole dataset, with colors indicating SNP calling parameters.
