## Supplementary figure 17 for "Extensive immune receptor repertoire diversity in disease-resistant rice landraces"

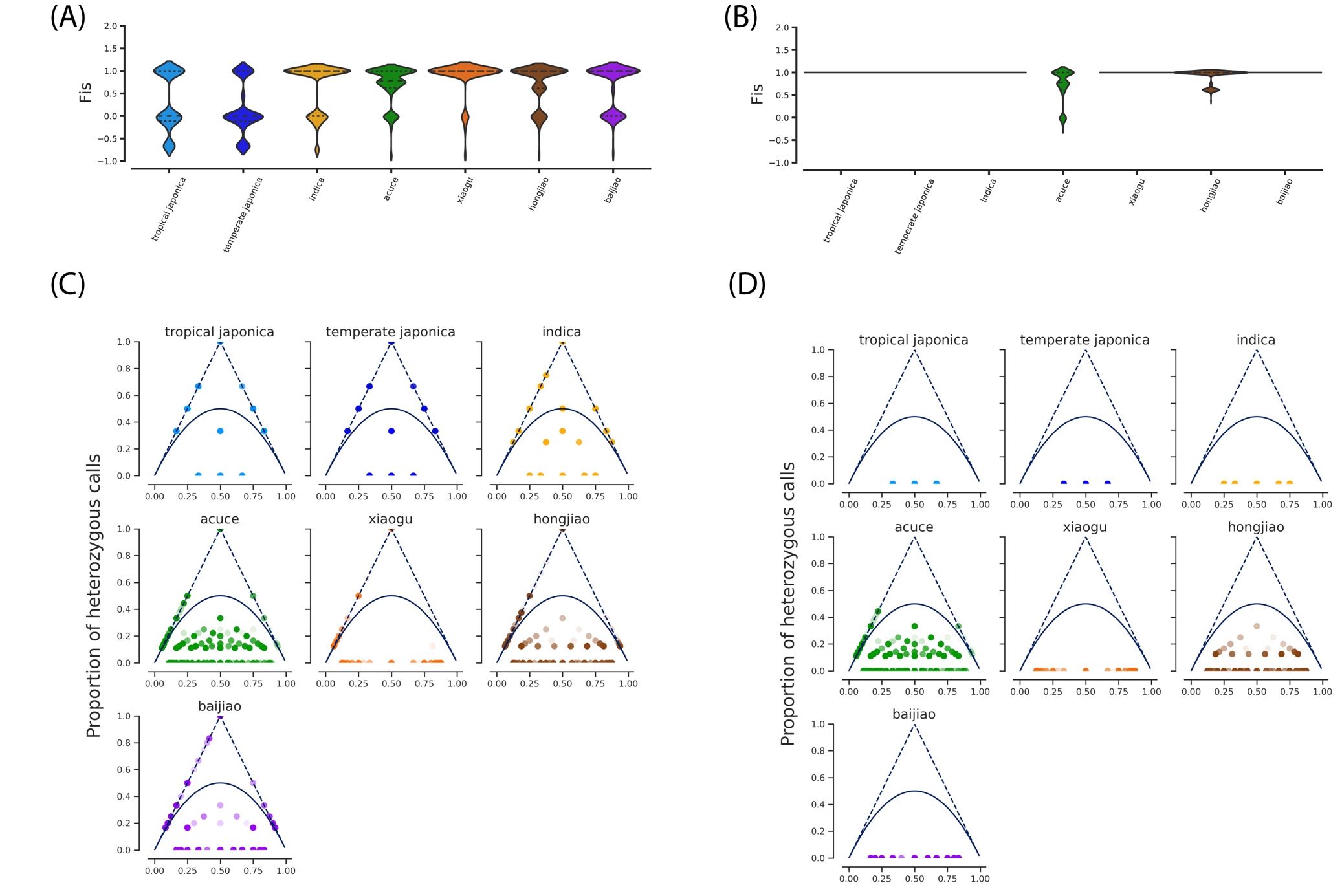
Supplementary Figure 17. SNP filtering. Distribution of the inbreeding coefficient Fis in groups of >2 accessions, before (A) and after (B) filtering for excess heterozygosity. Proportion of heterozygous calls versus frequency of non-reference allele in groups of >2 accessions, before (C) and after (D) filtering for excess heterozygosity. Solid lines: Hardy-Weinberg equilibrium. Dashed lines: maximum observed heterozygosity (proportion of heterozygous calls=2p, p being the minor allele frequency).
