## Supplementary figure 18 for "Extensive immune receptor repertoire diversity in disease-resistant rice landraces"

(A) GBS

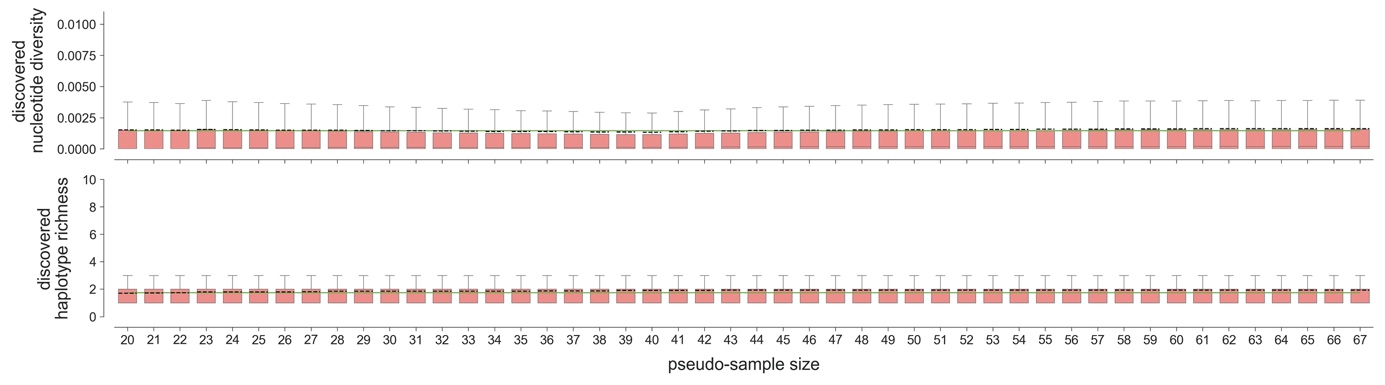

(B) RenSeq

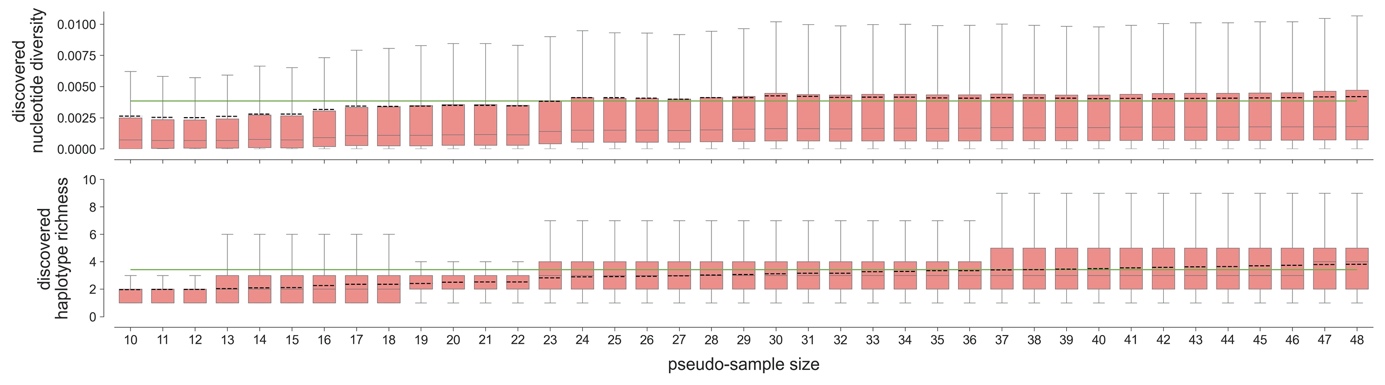

Supplementary Figure 18. Fraction of nucleotide diversity and haplotype richness recovered by subsampling datasets for (A) GBS loci excluding NLRs, (B) RenSeq data. Resampling was performed by taking 100 combinations of n accessions, n ranging from ten or twenty to sample size minus one (sample size is N=49 for RenSeq, N=68 for GBS). Solid horizontal green line is 0.9*max[average value at each pseudo-sample size]. In boxplots, dashed black line is mean, solid grey line is median.
