## Supplementary figure 19 for "Extensive immune receptor repertoire diversity in disease-resistant rice landraces"

(A) Acuce

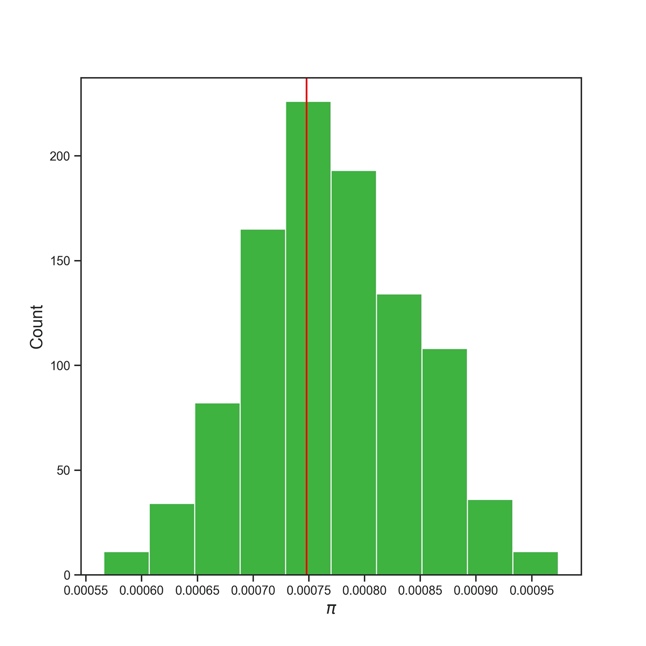

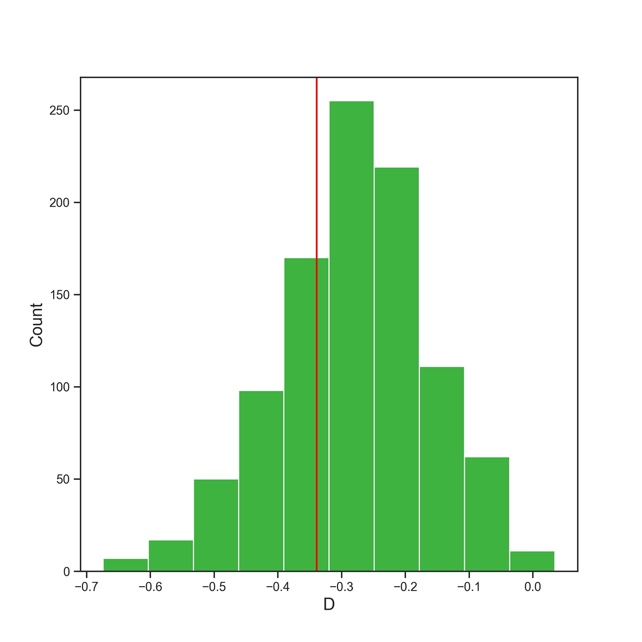

(B) Baijiao

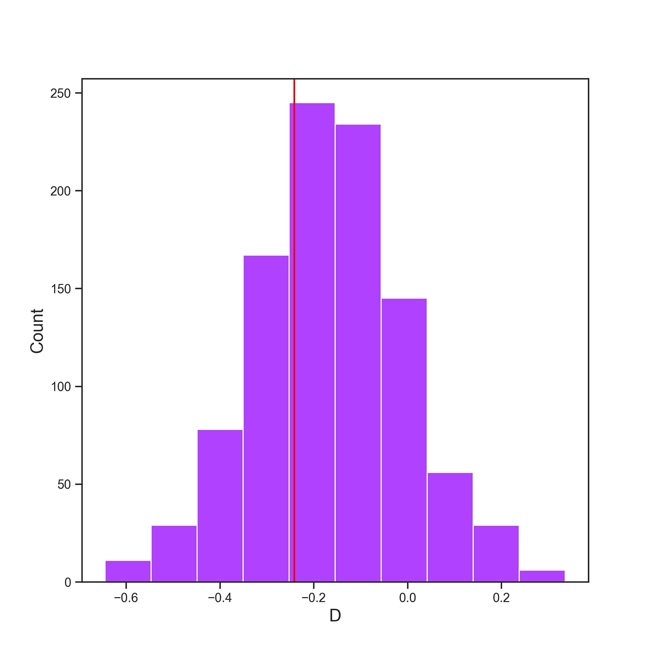

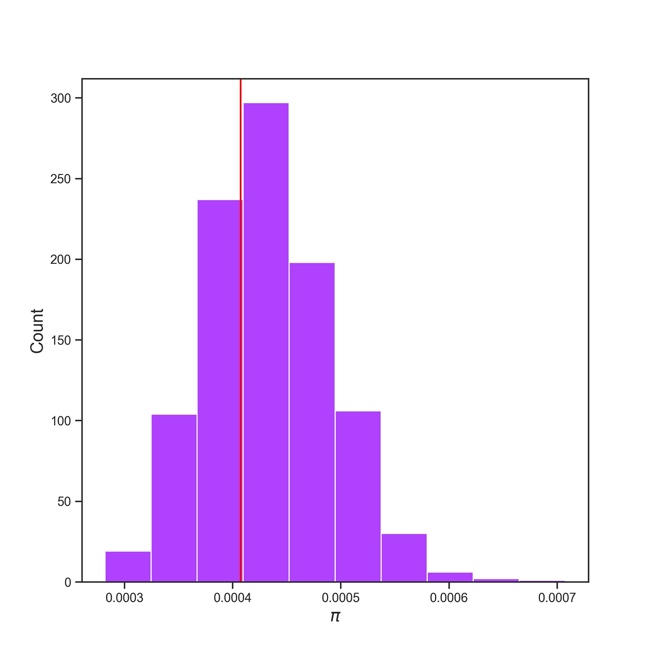

(C) Hongjiao

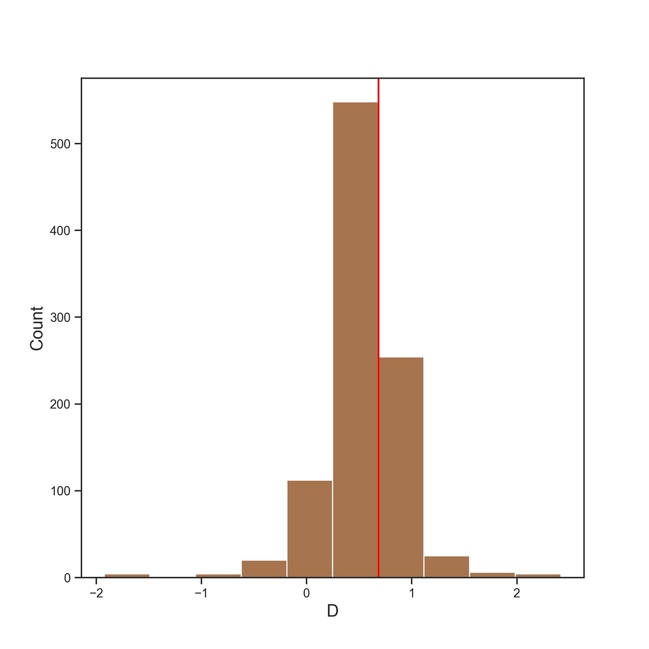

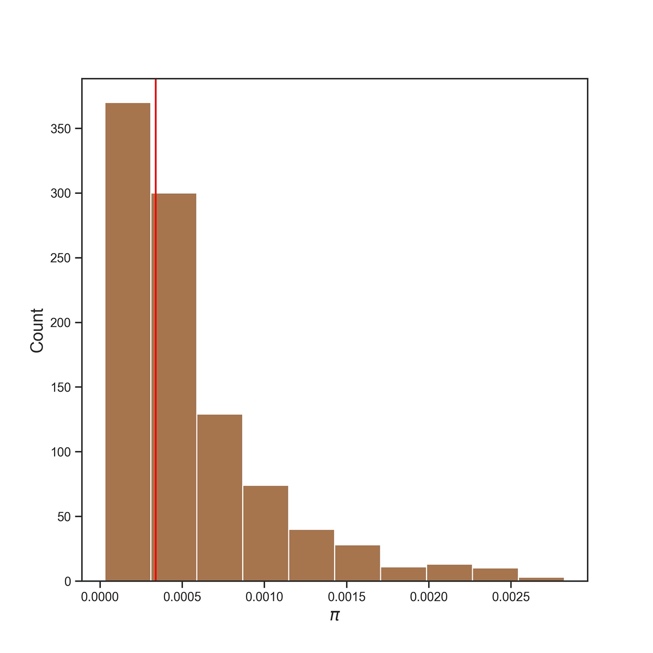

(D) Xiaogu

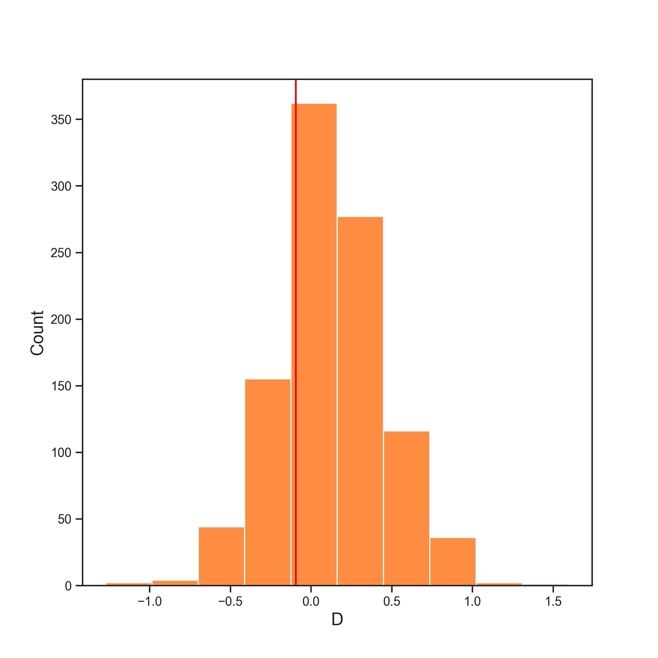

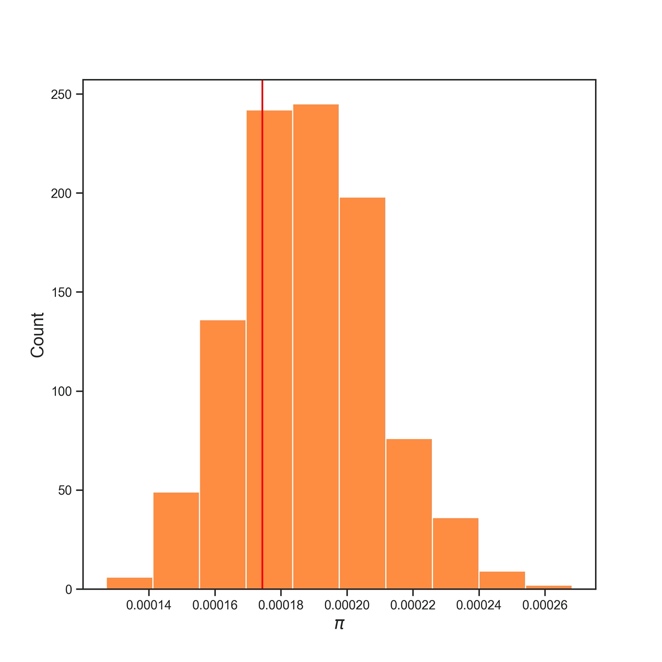

(E) indica landraces

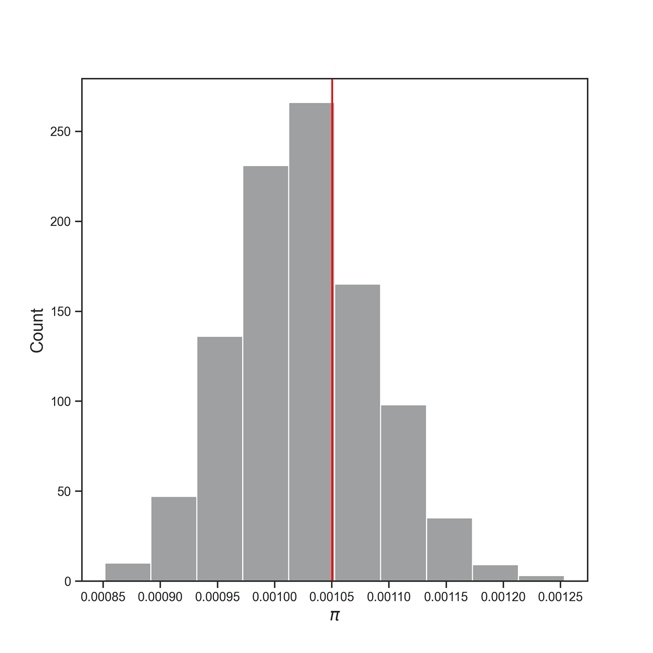

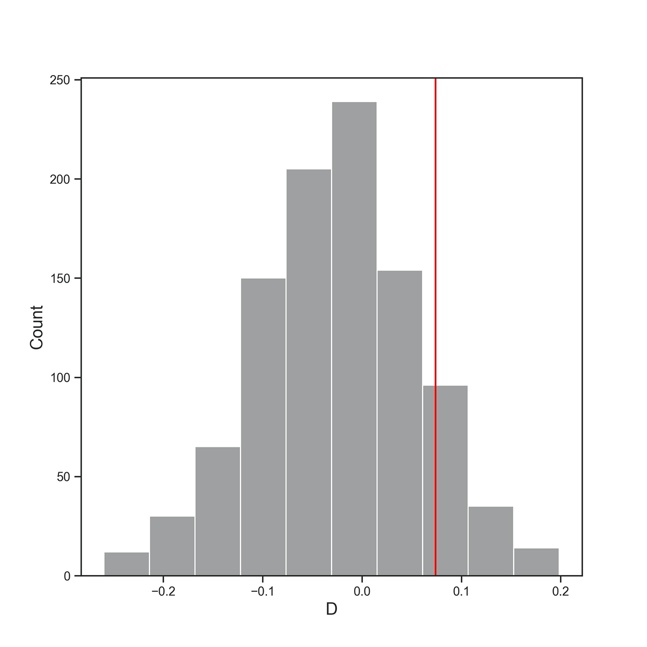

(F) indica

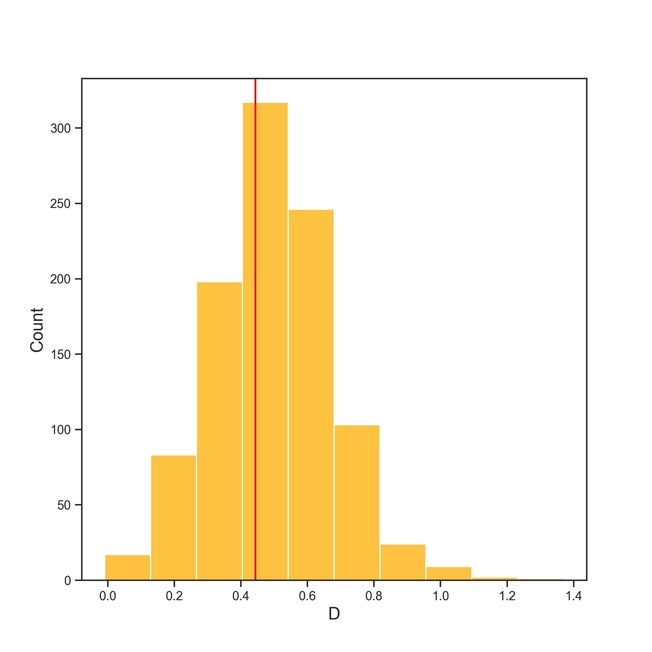

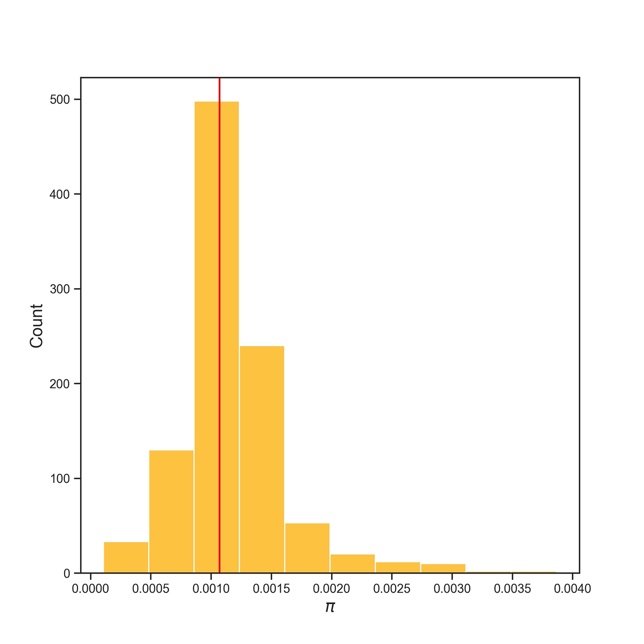

(G) japonica

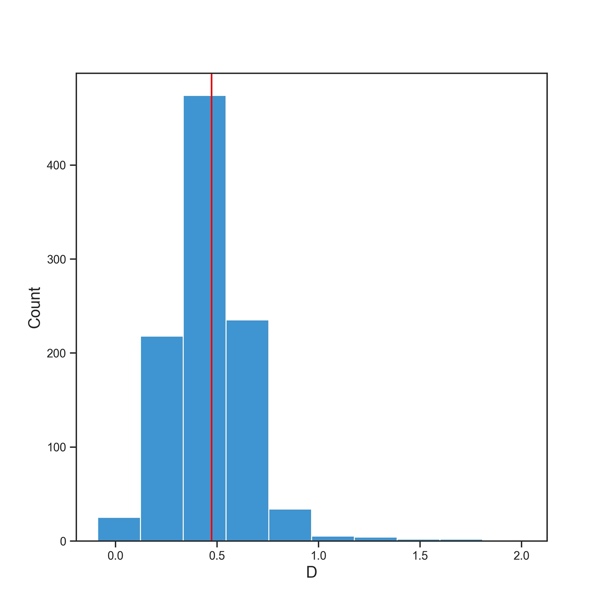

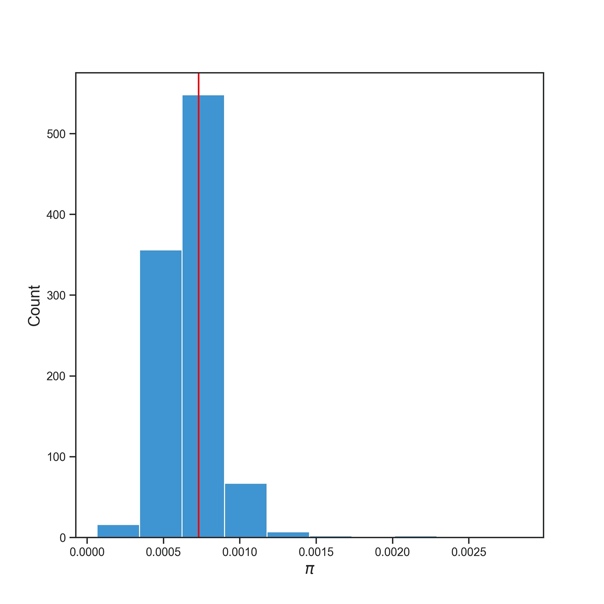

Supplementary figure 19. Results of posterior predictive checks^1^. Red lines indicate summary statistics (Tajima’s D and nucleotide diversity π) computed from observed datasets. Histograms represent summary statistics computed from datasets simulated by sampling multivariate parameters from posterior distributions (see details in Methods section).
