## Supplementary table 3 for "Extensive immune receptor repertoire diversity in disease-resistant rice landraces"

Supplementary Table 3. Posterior probabilities of the demographic models compared using approximate Bayesian computations.

| Population | Models | | | |
| --- | --- | --- | --- | --- |
|  | constant size | bottleneck | exponential growth | two epochs contraction |
| indica landraces | 0.6770 | 0.0066 | 0.1622 | 0.1541 |
| Acuce | 0.0036 | 0.0075 | 0.9889 | 0.0000 |
| Baijiao | 0.0644 | 0.2637 | 0.6689 | 0.0030 |
| Hongjiao | 0.0249 | 0.1591 | 0.0118 | 0.8042 |
| Xiaogu | 0.1249 | 0.6254 | 0.2326 | 0.0170 |
| indica | 0.0000 | 0.0000 | 0.0000 | 1.0000 |
| japonica | 0.0001 | 0.0016 | 0.0000 | 0.9983 |
