## Supplementary table 11 for "Extensive immune receptor repertoire diversity in disease-resistant rice landraces"

Supplementary Table 11. Prior distributions for demographic parameters in demographic models compared using approximate Bayesian computation. Parameters values were sampled from uniform distributions. Model parameters are described in the main text.

| Demographic model | Demographic parameters | Min. | Max. |
| --- | --- | --- | --- |
| Standard | *N1* | 10 | 100000 |
| Instantaneous bottleneck | *NI* | 10 | 20000 |
|  | *T1* | 0 | 1000 |
|  | *ST* | 0 | 200 |
| Exponential growth | N1 | 1000 | 100000 |
|  | N2 | 10 | N1 |
|  | T1 | 0 | 1000 |
|  | T2 | T1 | 20000 |
| Two epochs population expansion | N1 | 1000 | 100000 |
|  | N2 | 10 | 1000 |
|  | T | 10 | 10000 |
