## Supplementary table 12 for "Extensive immune receptor repertoire diversity in disease-resistant rice landraces"

Supplementary Table 12. Confusion matrix based on 100 samples for each demographic model. The simulations used here were carried out for the indica landrace group, but similar results were obtained for other populations and groups. Tolerance rate: 0.5%. This confusion matrix shows that, for instance, 91 out of the 100 datasets simulated under the bottleneck model were classified correctly.

| Population/group | Models | | | |
| --- | --- | --- | --- | --- |
|  | bottleneck | exponential growth | constant size | two epochs contraction |
| bottleneck | 91 | 5 | 0 | 4 |
| growth | 16 | 68 | 15 | 1 |
| Constant size | 15 | 11 | 74 | 0 |
| Two epochs contraction | 4 | 0 | 3 | 93 |
