## Supplementary table 13 for "Extensive immune receptor repertoire diversity in disease-resistant rice landraces"

Supplementary Table 13. Goodness-of-fit analyses for approximate Bayesian computations. The test statistic is the median of the distance between accepted summary statistics and the observed summary statistics. The null distribution is estimated using already performed simulations as pseudo-observed datasets. The p-value is computed as the proportion of test statistics obtained under a given model that are larger than the observed test statistic. One hundred pseudo-observed datasets were used per model.

| Population/group | Models | | | |
| --- | --- | --- | --- | --- |
|  | constant size | bottleneck | exponential growth | two epochs contraction |
| Acuce | 0.02 | 0.11 | 0.84 | 0.01 |
| Baijiao | 0.04 | 0.13 | 0.69 | 0.04 |
| Hongjiao | 0.01 | 0.07 | 0.04 | 0.54 |
| Xiaogu | 0.17 | 0.25 | 0.44 | 0.04 |
| indica landraces | 0.32 | 0.09 | 0.41 | 0.24 |
| indica | 0.00 | 0.06 | 0.04 | 0.83 |
| japonica | 0.01 | 0.07 | 0.05 | 0.64 |
